# Plasticity, not genetics, shapes individual responses to thermal stress in non-native populations of the European green crab (*Carcinus maenas*)

**DOI:** 10.1101/2025.04.22.650023

**Authors:** Yaamini R. Venkataraman, Julia C. Kelso, Catlin Payne, Heidi L. Freitas, Jasmine Kohler, Carolyn K. Tepolt

## Abstract

Temperature is a major driver of individual performance in ectotherms, with this impact depending on stressor intensity and duration. Differences in individual response across temperature, time, and populations are shaped by the interplay between evolutionary adaptation and phenotypic plasticity. Some populations are able to thrive in novel and changing environments despite limited genetic diversity, raising the question of how plasticity and adaptation interact after significant genetic diversity loss. The European green crab (*Carcinus maenas*) is a textbook example of this phenomenon: invasive populations boast a broad thermal tolerance and exceptional thermal flexibility even after repeated genetic bottlenecks. Despite this loss of diversity overall, prior work has found a strong population-level association between variation at a specific extended genomic region (supergene), cold tolerance, and sea surface temperature. We conducted a series of three experiments using righting response to characterize sublethal thermal tolerance and plasticity in introduced green crab populations, then determined if these factors were associated with supergene genotype for individual adult crabs. Crabs showed signs of stress after exposure to a 30°C heat shock in one experiment. Interestingly, a second experiment exposing *C. maenas* to repeated 24-hour heat shocks showed that prior heat shock conferred beneficial plasticity during a subsequent event. The third experiment examined cold acclimation over multiple timepoints up to 94 hours. At 5°C, certain crabs exhibited an acclimatory response where righting slowed dramatically at first, and then gradually sped up after a longer period of cold exposure. Several crabs failed to right at 1.5°C, which could be indicative of dormancy employed to reduce energy consumption in colder conditions. There were no significant relationships between individual plasticity and supergene genotype in any experiment. Linking population-level genetic associations with individual-level physiology is complex, and reflects the impact of environmental conditions such as temperature throughout life history in shaping adult phenotype. Our results highlight the robust thermal tolerance and plasticity that adult green crabs maintain despite a substantial reduction in genetic diversity, and underscore the importance of probing population-level genotype-phenotype associations at the individual level.

## Introduction

Temperature is a primary driver of physiology in marine invertebrates (Somero, Lockwood and Tomanek 2017), and substantial effort has focused on quantifying its impacts on these species (Yao and Somero 2014). In some circumstances, acute thermal stress may elicit a pronounced stress response, such as higher mortality, which then subsides after extended periods of stress exposure (Zou *et al*. 2023). Alternatively, short-term exposure can be less stressful due to acclimatory potential not present in response to longer-term stressors (Madeira *et al*. 2018; Morley *et al*. 2019). Populations within a species can also exhibit distinct responses to thermal stress due local adaptation or thermal history (Sanford and Kelly 2011; Fields, Cox and Karch 2012; Falfushynska, Phan and Sokolova 2016). These different responses across temperature, time, and populations can be attributed to various mechanisms, including shifts in metabolic pathways, changes in lipid membrane composition, and differential gene expression (Pörtner 2002; Somero 2010, 2012). Understanding these mechanisms may aid in predicting organismal responses to more frequent thermal extremes (Calvin *et al*. 2023) and uncover why certain species and populations thrive in inherently stressful conditions (Somero 2010).

Intraspecific variation in physiology often arises from some combination of evolutionary adaptation and phenotypic plasticity (Fox *et al*. 2019). Evolutionary adaptation depends on genetic differences within a species that alter fitness in a given environment (Fox *et al*. 2019). For example, genetic differences over five decades in *Daphnia magna* were associated with changes in critical thermal maxima (Cuenca Cambronero *et al*. 2018). Phenotypic plasticity, or the ability for a single genotype to produce more than one phenotype, also plays a critical role in shaping an individual organism’s thermal tolerance (Hendry 2016; Fox *et al*. 2019). Thermal priming is a common plastic response in many species including marine invertebrates, in which prior exposure to heat stress confers beneficial plasticity in response to future shocks (Martell 2022; Glass *et al*. 2023; Abbas *et al*. 2024). Repeated heat shocks boosted thermal tolerance of the abalone *Haliotis tuberculata* with no long-term detrimental effects (Abbas *et al*. 2024), while prior exposure to elevated temperatures improved *Nematostella vectensis* growth, development, and acute heat tolerance (Glass *et al*. 2023). Populations in variable environments often evolve greater capacity for plasticity (Handelsman *et al*. 2013; Landy *et al*. 2020). However, acclimatory plasticity may not fully mediate negative impacts of thermal stress (Gunderson and Stillman 2015), and tradeoffs between tolerance and plasticity can produce thermally-tolerant genotypes with lower plasticity (Barley *et al*. 2021; Griffiths *et al*. 2024). For example, experimental evolution of the copepod *Acartia tonsa* produced genotypes with a 2-5°C increase in LD50 over 40 generations, but with decreases in plasticity of over 60% (Sasaki and Dam 2021). Understanding the interplay of these mechanisms may uncover why certain species thrive in inherently stressful conditions (Somero 2010).

If changes in thermal physiology are driven by selection and plasticity, then does plasticity alone explain why some populations boast broad environmental tolerances with limited genetic diversity? Several introduced species thrive in novel environments even after substantial losses of genetic variation (Roman and Darling 2007; Estoup *et al*. 2016). However, even highly bottlenecked populations often retain some genetic diversity, including (and perhaps especially) at adaptive markers (Swindell and Bouzat 2005; Oliver and Piertney 2012; Teixeira and Huber 2021). Population genomic research has identified specific genetic markers correlated with various environmental parameters across populations in many introduced species (Schwander, Libbrecht and Keller 2014; Zhang *et al*. 2023; Ma *et al*. 2024), but it remains a challenge to link these population-level association studies with individual-level physiological mechanisms (Riginos *et al*. 2016; Lind and Lotterhos 2024). Some studies have successfully identified genome-phenome linkages at various levels of biological hierarchy. For example, genetic markers associated with warmer and more arid conditions are positively correlated with *Anopheles gambiae* larval thermal tolerance (Rocca *et al*. 2009), and flies (*Drosophila suboscura*) with putatively warm-adapted genotypes had higher abundance of heat shock proteins than flies with cold-adapted genotypes (Calabria *et al*. 2012). Individual survival within one population of adult dwarf surf clams *Mulina lateralis* at 30°C for 17 days was associated with variation in genes in the ETHR/EHF signaling pathway (Wang *et al*. 2024). Additionally, these genes were more highly expressed in heat-tolerant clams, further illustrating the importance of these population-level genetic markers to individual thermal tolerance (Wang *et al*. 2024).

The European green crab (*Carcinus maenas*) provides the perfect opportunity to study these interactions in genetically depauperate yet highly plastic populations. Like many non-indigenous species, *C. maenas* boasts broad thermal tolerance and plasticity (Davidson, Jennions and Nicotra 2011; Zerebecki and Sorte 2011; Sorte *et al*. 2013), with upper thermal limits higher and lower thermal limits lower than other temperate crustaceans (Tepolt and Somero 2014; Tepolt 2024). Their evolution in variable estuarine environments spanning 4-23°C in the native range may have shaped this wide thermal optima range (Compton, Leathwick and Inglis 2010; Jost, Podolski and Frederich 2012). A variety of mechanisms may underpin this broad thermal tolerance, including “dumping” hemocyanin-bound oxygen to extend upper thermal limits (Jost, Podolski and Frederich 2012) and altering hemolymph magnesium content in colder temperatures to reduce the blockage of calcium channels that otherwise would impair movement (Frederich *et al*. 2000; Wittmann *et al*. 2010). Seasonal differences in thermal limits (Cuculescu, Hyde and Bowler 1998; Hopkin *et al*. 2006) and rapid homeoviscous adaptation (Chapelle 1978, 1986; El Babili *et al*. 1997) observed in *C. maenas* underscore the importance of phenotypic plasticity to the species’ success across broad thermal gradients.

Green crabs in invasive North American populations are characterized by substantial losses of genome-wide genetic diversity, but still maintain broad thermal plasticity (Tepolt and Palumbi 2015) (see **Appendix S1** for a detailed *C. maenas* invasion history). North American *C. maenas* inhabit a broader thermal range than in their native range (Compton, Leathwick and Inglis 2010), and are thriving in sub-arctic waters in Newfoundland (Kelley, de Rivera and Buckley 2013; Best, McKenzie and Couturier 2017). While introduced *C. maenas* lost substantial genetic diversity, the variation remaining in these populations may underpin thermal plasticity. Population genomics of *C. maenas* revealed a group of single nucleotide polymorphisms (SNPs) in strong linkage disequilibrium that were significantly associated with thermal tolerance on a population level; these SNPs likely represent an extended region of reduced recombination, or supergene (Tepolt and Palumbi 2020). Crab populations with higher heart rates at 0°C had a higher frequency of the putatively “cold-adapted” supergene allele (Tepolt and Palumbi 2020), while populations experiencing warmer average sea surface temperatures had a higher frequency of the putatively “warm-adapted” supergene allele (Tepolt and Palumbi 2020). In a separate study, this supergene represented nearly all of the genomic variation associated with sea surface temperature in northeast Pacific populations (Tepolt *et al*. 2022). Among the dozens of single-nucleotide differences between the two main supergene alleles are two nonsynonymous SNPs in *hypoxia-inducible factor 1-alpha* (Tepolt and Palumbi 2020): as a key regulator of hypoxia, it may also be essential in marine invertebrate responses to temperature and offers a potential functional link between supergene genotype and thermal phenotype (Pörtner, Langenbuch and Michaelidis 2005; Pörtner, Bock and Mark 2017). While the genomic data suggest a strong link between genomic adaptation and population-level thermal tolerance, it is unclear if or how these differences translate to individual thermal plasticity. Supergene genotype had no impact on individual response to acute cold stress in adult *C. maenas* from several northwest Atlantic populations (Coyle *et al*. 2019); however, the crabs were only exposed to 4.5°C for a maximum of six hours. Given the complexities of linking population-level associations with individual physiology (Gagnaire and Gaggiotti 2016; Riginos *et al*. 2016; Tam *et al*. 2019; Lind and Lotterhos 2024), investigating the role of the supergene at the individual level may help elucidate the mechanisms that shape thermal tolerance in the species.

We conducted three experiments to investigate individual thermal tolerance and plasticity in adult crabs in response to thermal extremes, then tested whether the observed responses were linked to the temperature-associated supergene (**Figure 1**). We first compared responses to acute heat shock in crabs from Washington and Maine (“Experiment 1”), and hypothesized that crabs from Maine, which experience higher summer temperatures, would display a higher heat tolerance than their Washington counterparts. To understand plasticity in response to heat stress, we exposed crabs from Washington to repeated acute heat shocks (“Experiment 2”). We predicted that “priming” from previous heat shock exposure would confer beneficial plasticity in response to subsequent acute heat stress. Finally, we characterized cold tolerance and plasticity for crabs from Massachusetts and Washington over multiple timepoints across 94 hours (“Experiment 3”), to explore the extent and time scale of plasticity at low temperatures. We expected that crabs would slow upon initial cold exposure, but might exhibit evidence of thermal acclimation over longer periods. We also expected crabs from MA to have a higher cold tolerance than WA crabs due to lower winter temperatures in the northwest Atlantic than in the northeast Pacific. Across all three experiments, we tested whether and how supergene genotype influenced thermal tolerance and plasticity. We expected crabs with at least one copy of the putatively warm-adapted allele to be more robust to heat stress. Conversely, we hypothesized that the putatively cold-adapted allele would improve response to short- and medium-term cold acclimation. Our results show how investigating physiological response at the individual organism level can elucidate how plasticity and genetic background shape thermal tolerance in genetically depauperate but highly successful populations.

**Figure 1.**
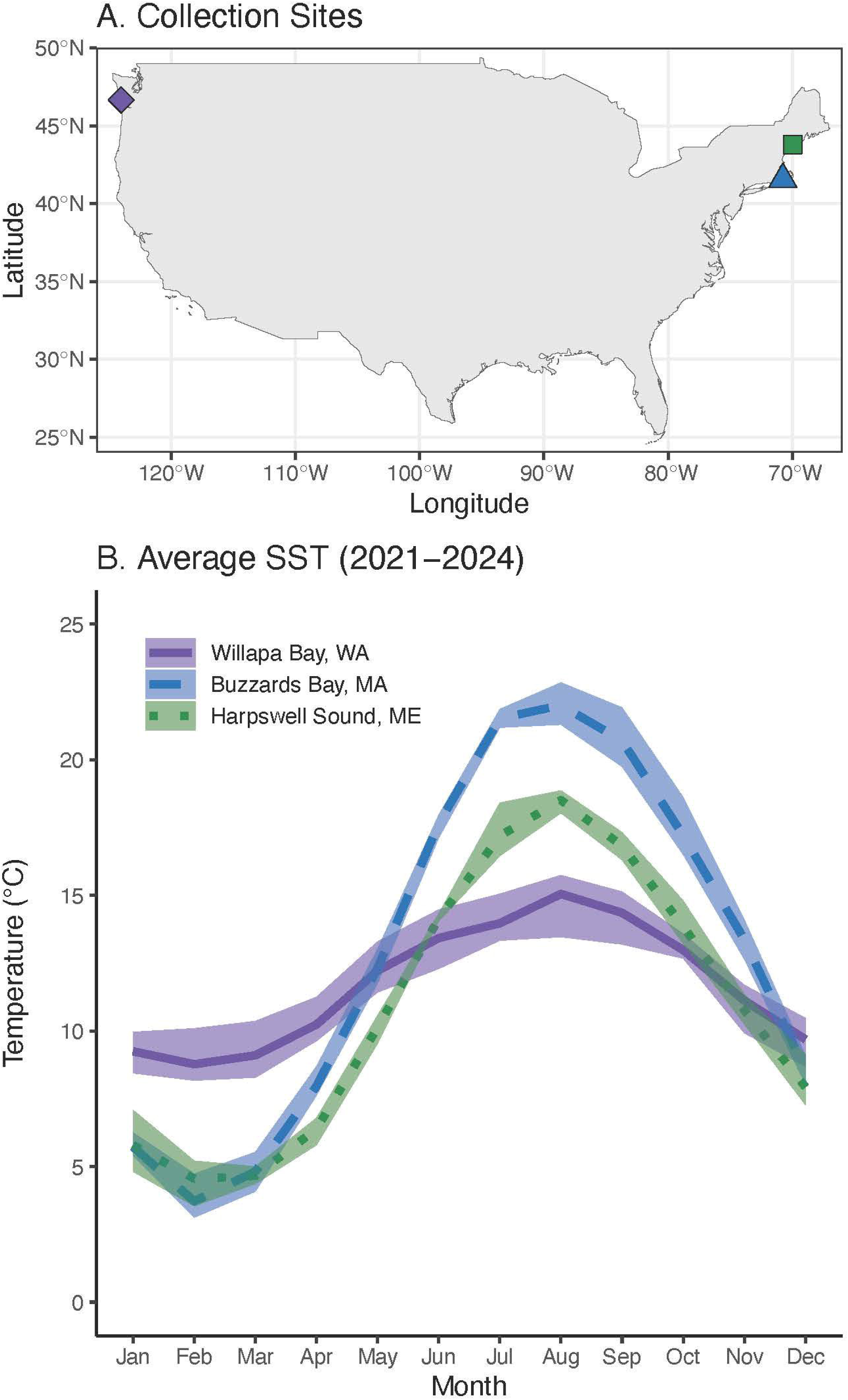
**A**) Green crab collection sites for experiments (purple diamond = Willapa Bay, WA; blue triangle = Buzzards Bay, MA; and green square = Harpswell Sound, ME). **B**) Average sea surface temperature (SST) for 2021-2024 near collection sites (solid purple line = Willapa Bay, WA; long blue dashes = Buzzards Bay, MA; small green dashes = Harpswell Sound, ME). SST data were obtained from NOAA’s OI SST V2 High Resolution Dataset provided by the NOAA PSL, Boulder, Colorado, USA, from their website at https://psl.noaa.gov (Huang *et al*. 2021). A four-year mean, SST minimum, and SST maximum were calculated for each month. Solid lines represent four-year averages, while ribbons represent ranges.

## Methods

### General experimental setup

A series of experiments was conducted to assess the impact of short-(∼24 hours) to medium-term (94 hours) heat and cold stress on introduced *C. maenas* populations in both the northwest Atlantic and northeast Pacific. Experimental crabs were individually labeled using waterproof paper and superglue, and the propus and dactyl segments from the third walking leg on the right side (dorsal view) were preserved in 95% ethanol for subsequent genotyping. These joints were taken as the third walking leg has no significant role in crab righting behavior (Young, Peck and Matheson 2006). If the crab did not have a third walking leg on the right side, the third walking leg was taken from the left.

Crabs were placed in 21 cm x 26 cm x 41 cm (22.386 L) tanks filled with 800 micron filtered raw seawater from Great Harbor, MA, USA (Experiments 1 and 3) or 50 micron filtered raw seawater from Vineyard Sound, MA, USA (Experiment 2), and equipped with an air stone and filter (**Appendix S2**). Crabs were distributed to balance sex, carapace width, and integument color among tanks. Crabs with orange-red or red integument color were not included in experiments; these colors indicate an extended time in intermolt and “red” green crabs have less robust tolerance to a range of physiological stressors (Styrishave, Rewitz and Andersen 2004). Remaining crabs had integument colors at the beginning of the experiment ranging from blue to yellow-orange. Tanks were checked for mortalities, cleaned, and fed daily (**Appendix S2**). Ammonia concentrations were checked every third day and filters were replaced every five days (**Appendix S2**). A 12-hour:12-hour light:dark cycle was maintained for all tanks (Tepolt and Somero 2014).

Crabs were initially kept in acclimation conditions for at least five days (see details below). Water temperature was maintained during the acclimation period using either a flowing water bath or environmental chamber manipulation, and heated or cooled using a 300-Watt Deluxe Titanium Heating Tubes (Finnex, USA) with an external temperature controller (bayite, China) or environmental chamber, respectively (**Appendix S2**). Water temperature in each tank was recorded every 15 minutes using two HOBO Data Loggers (Onset, USA), one at the water surface and one at the bottom of the tank. A non-parametric Kruskal-Wallis test (`kruskal.test` from R `stats` package (R Core Team 2024) was used to confirm that temperature treatments were statistically different from each other and that replicate tanks were not different from each other.

### Measuring thermal tolerance with time-to-right

Time-to-right (TTR) was used in all experiments to assess sublethal thermal tolerance. TTR was defined as the number of seconds *C. maenas* spent on righting behavior (defined in (Young, Peck and Matheson 2006)), including “hesitation time” after crabs are placed dorsal side up but before moving their fifth pereopod pair. Individual crabs were placed with their dorsal side on the bottom surface of the tank and given 90 seconds to right. If a crab did not right within 90 seconds, it was manually righted and recorded as “not righting” for that trial. Crabs were rested for 10 seconds before repeating the TTR trial twice for a total of three TTR measurements per crab per trial. Readings were always conducted by the same person within an experiment to reduce error. To control for other factors that may influence righting response, crabs were weighed, measured, sexed, and assessed for integument color and missing legs after each TTR trial.

### Supergene genotyping

Crabs were genotyped at a single-nucleotide polymorphism (SNP) in the *structural maintenance of chromosomes protein 3* (SMC) gene to understand how individual variation at the putative supergene impacts thermal tolerance. This SNP is part of a group of SNPs in very strong linkage disequilibrium in the putative supergene, and is diagnostic of supergene allele (Tepolt & Palumbi 2020). DNA was extracted from ∼4 mm^2^ of leg joint tissue using the DNeasy Blood & Tissue Kit (QIAGEN, Valencia, CA, USA) following manufacturer’s instructions. Incubation with the ATL Lysis Buffer and Proteinase K occurred for at least two hours. The final eluate was pipetted back onto the spin column, incubated, and re-eluted to increase DNA yield.

Genotyping was done with one of two equivalent methods. For Experiment 2, a 126 bp region of the SMC gene containing a C/T single nucleotide polymorphism (SNP) diagnostic for supergene allele was amplified using the SMC F and R primers and methods specified in Coyle et al. (2019) (**Table S1**). Amplified product was sent to Sequegen (Worcester, MA, USA) for Sanger sequencing. Resultant chromatograms were edited and aligned using Sequencher v.5.4.6 (Gene Codes, Ann Arbor, MI, USA). Samples from Experiments 1 and 3 were genotyped using a restriction digest assay. A 690 bp region of the SMC gene containing the same diagnostic C/T SNP was amplified using SMC_long F and R primers (**Table S1**). Amplified product was digested with 0.5 µL of Alul enzyme (Promega, USA or ThermoFisher, USA) at 37°C for one hour. The recognition site includes a CG dinucleotide at 88 bp in the PCR product, meaning that digestion only occurs for C alleles There is an additional non-target cut site at 477 bp that is digested for all genotypes. Digested product was run out on a 1.5% agarose gel with GelRed® Nucleic Acid Gel Stain (Biotium, Inc, Fremont, CA, USA) at 90 V for at least 65 minutes then imaged. A 477 bp band indicated TT homozygotes, a 389 bp band indicated CC homozygotes, and both bands indicated CT heterozygotes (**Figure S1**). Both CC and CT genotypes had also an 88 bp band. All genotypes had a 213 bp band due to digestion at the non-target site (**Figure S1**). The C allele is found in the cold-adapted supergene allele, while the T allele is found in the warm-adapted supergene allele.

### Experiment 1: Responses to acute heat shock across populations

This experiment gauged *C. maenas* response to an acute heat shock in WA and ME populations (**Appendix S3**). The WA and ME crabs were placed in tanks with 15-17 crabs each, separated by site. All crabs were acclimated at room temperature (∼22-24°C) for at least five days. After the acclimation period, crabs were exposed to a heat ramp of ∼1°C/hour to 30°C over 6 hours, then remained at 30°C for 22 hours (**Figure S3**). TTR was assessed before and at the end of the 30°C heat shock. Prior to analysis, the three replicate TTR measurements for each individual crab at each time point were averaged and log-transformed. Trials where the crab did not right itself within 90 seconds were excluded from averaging. After experiments were concluded, crabs were sacrificed and dissected to identify and count trematode parasite cysts in the hepatopancreas tissue per Blakeslee et al. (2015), with the modification that all hepatopancreas tissue was examined on 12.5 cm x 7.5 cm squash plates. Parasite intensity was calculated by dividing metacercarial cyst count by crab carapace width. Since *C. maenas* populations in the northwest Atlantic are parasitized while counterparts in the northeast Pacific are not, parasite intensity was quantified to control for any extraneous factors impacting righting when comparing ME and WA crab responses.

Linear mixed effects models (`lmer` from R `lme4` package; (Bates *et al*. 2015)) were used in three steps to evaluate how various factors impacted log-transformed average righting response time with crab ID used as a random effect to correct for repeated measurements of the same animals. First, a null model was constructed with crab sex, integument color, carapace width, weight, whether or not a crab was missing a fifth pereopod (“swimmer”), and parasite intensity (ME only) as explanatory variables to understand how these factors explained variation in righting response. An Analysis of Variance (ANOVA; `anova` from R `stats` package (R Core Team 2024)) was used to perform a likelihood ratio test and compare the null model with a model testing the influence of a specific factor (*ie*. only carapace width or parasite infection intensity). Second, any significant variables identified by ANOVA tests were included in a new model along with timepoint. Timepoint was modeled as a categorical variable because it directly corresponded with heat shock (hour 0 = no heat shock, hour 22 = heat shock). Significant factors (*ie*. timepoint, any significant demographic variables) identified by ANOVA tests were then included in a third and final null model. This “no genotype” null model was compared to three separate models containing significant factors identified by the previous two null models and either genotype, presence of the C allele, or presence of the T allele. A final set of ANOVA tests were used to discern if these genotype-specific factors significantly impacted log-transformed average righting response. For all models, *P*-values were corrected for multiple comparisons using a Bonferroni correction. Factors were considered significant when adjusted p < 0.05. Normality and homoscedasticity were verified visually. Pairwise tests were conducted with `emmeans` v.1.11.0 for any significant factor (Lenth *et al*. 2018).

### Experiment 2: The role of “heat priming” in shaping thermal tolerance

The goal of this experiment was to understand how repeated heat shocks impacted *C. maenas* thermal plasticity. After the acclimation period (**Appendix S4**), two tanks (treatment group) were exposed to a ∼30°C heat shock for 20 hours, while the other two tanks (control group) were maintained at 15°C. After the initial heat shock, all tanks were maintained at 15°C for 48 hours. Both the control and treatment groups were then exposed to a second heat shock of ∼30°C for 24 hours.

Righting response was measured for all crabs after the acclimation period (day 0), at the end of the first heat shock (day 1), and at the end of the second heat shock (day 4). Measurements taken for crabs experiencing heat shocks were conducted at 30°C. A three-step series of linear mixed effects models was used to assess the impact of sex, integument color, carapace width, weight, a missing swimmer, treatment, day, the interaction of treatment and day, genotype, presence of the C allele, and presence of the T allele on log-transformed average TTR, with crab ID used as a random effect to correct for repeated measurements of the same animals. (see *Experiment 1* for specific methods). For this experiment, day was modeled as a categorical predictor, as it was a proxy for heat shock timing. Pairwise tests were conducted with `emmeans` v.1.11.0 for any significant factor (Lenth *et al*. 2018). Oxygen consumption was also measured for a subset of crabs in this experiment (see **Appendix S4** for details).

### Experiment 3: Changing cold tolerance across populations over time

This experiment investigated how short- and medium-term exposure to cold temperatures influenced thermal tolerance and plasticity in green crabs from MA and WA over the course of 94 hours (**Appendix S5**). At hours 0, 4, 22, 28, 46, and 94, righting response was assessed for at least seven crabs per tank. A series of linear mixed effects models was used to assess the impact of sex, integument color, carapace width, weight, a missing swimmer, treatment, time, the interaction of treatment and time, genotype, presence of the C allele, and presence of the T allele on log-transformed average TTR, with crab ID used as a random effect to correct for repeated measurements of the same animals (see *Experiment 1* for specific methods). For this experiment, timepoint was modeled as a continuous predictor. Pairwise tests were conducted with `emmeans` v.1.11.0 for any significant factor (Lenth *et al*. 2018). To obtain pairwise differences between different timepoints, timepoint was re-estimated as a categorical variable.

A different set of models was used to evaluate how various factors impacted failure to right within the 90 second cutoff. For these models, TTR for each crab was coded as a “success” (righting within 90 seconds) or “failure” (no righting within 90 seconds). Binomial linear mixed effects models with crab ID as a random effect were used to understand how various factors impacted binary righting response in three steps. First, a null model was constructed with crab sex, integument color, carapace width, weight, and whether or not a crab was missing a swimmer as explanatory variables to understand how these factors explained variation in binary righting response. An Analysis of Variance (ANOVA; `anova` from R `stats` package (R Core Team 2024)) was used to perform a likelihood ratio test and compare the null model with a model testing the influence of a specific factor (*ie*. only carapace width or weight). Second, any significant variables identified by ANOVA tests were included in a new model along with temperature, timepoint, and their interaction. Significant factors (*ie*. timepoint, any significant demographic variables) identified by ANOVA tests were then included in a third and final null model. This “no genotype” null model was compared to three separate models containing significant factors identified by the previous two null models, and either genotype, presence of the C allele, or presence of the T allele. A final set of ANOVA tests were used to discern if these genotype-specific factors significantly impacted log-transformed average righting response. Factors were considered significant at p < 0.05 after Bonferroni correction.

Sample size decreased over the course of the experiment due to 5-6 crabs per treatment (1-2 crabs per tank) being randomly selected for dissection at each timepoint for future analyses. To account for this, a sensitivity analysis was used to confirm the validity of the average time-to-right and failure-to-right modeling results (see **Appendix S5** for details).

To compare responses to temperature between populations, separate linear mixed effects models were created to evaluate the impact of population and significant variables from the population-specific models on average time to right, or on failure to right. Generalized linear models (`glm` from R `stats` package (R Core Team 2024)) were also used to compare the effect of population on failure to right at each timepoint for crabs in the colder temperature treatment. All *P*-values were corrected for multiple comparisons using a Bonferroni correction.

## Results

### Experiment 1: Responses to acute heat shock across populations

Righting response was evaluated before and after a 22 hour heat shock to understand the factors that impact short-term heat tolerance. Timepoint had a significant impact on righting response in both populations (**Figure 2A**; WA comparison: X^2^ = 20.85, *P*-value < 0.0001; ME comparison: X^2^ = 19.17, *P*-value < 0.0001). Crabs from both WA and ME had significantly slower TTR after heat shock (mean_WA, 22 hours_ = 3.22 s ± 4.22 s; mean_ME, 22 hours_ = 2.22 s ± 2.15 s) than before (mean_WA, 0 hours_ = 1.16 s ± 0.26 s; mean_ME, 0 hours_ = 1.23 s ± 1.15 s). Neither genotype χ = 0.48, *P*-value = 1), presence of the C allele (χ = 3.02 × 10, *P*-value = 1), nor presence of the T allele (X^2^ = 3.24, *P*-value = 1) had a significant impact on average TTR in WA *C. maenas* (**Figure 2B**). A similar trend was identified in the ME population, which only had CT and CC crabs (**Figure S2B**). After heat shock, CT crabs had a slightly faster righting response (mean_ME, 22 hours, CT_ = 1.46 s ± 0.20 s) than CC crabs (mean_ME, 22 hours, CC_ = 2.67 s ± 2.62 s), but this difference was not significant after multiple test correction (genotype: X^2^ = 5.65, P-value = 0.12) (**Figure 2B**). Righting response was not impacted by sex, integument color, carapace width, weight, or whether or not a crab was missing a fifth pereopod in this or any other experiment (**Table S3**). Parasite infection intensity did not impact righting response in ME crabs (**Table S3**). Crab ID explained 4.11% and 21.69% of the variation in final WA and ME models (timepoint only), respectively.

**Figure 2.**
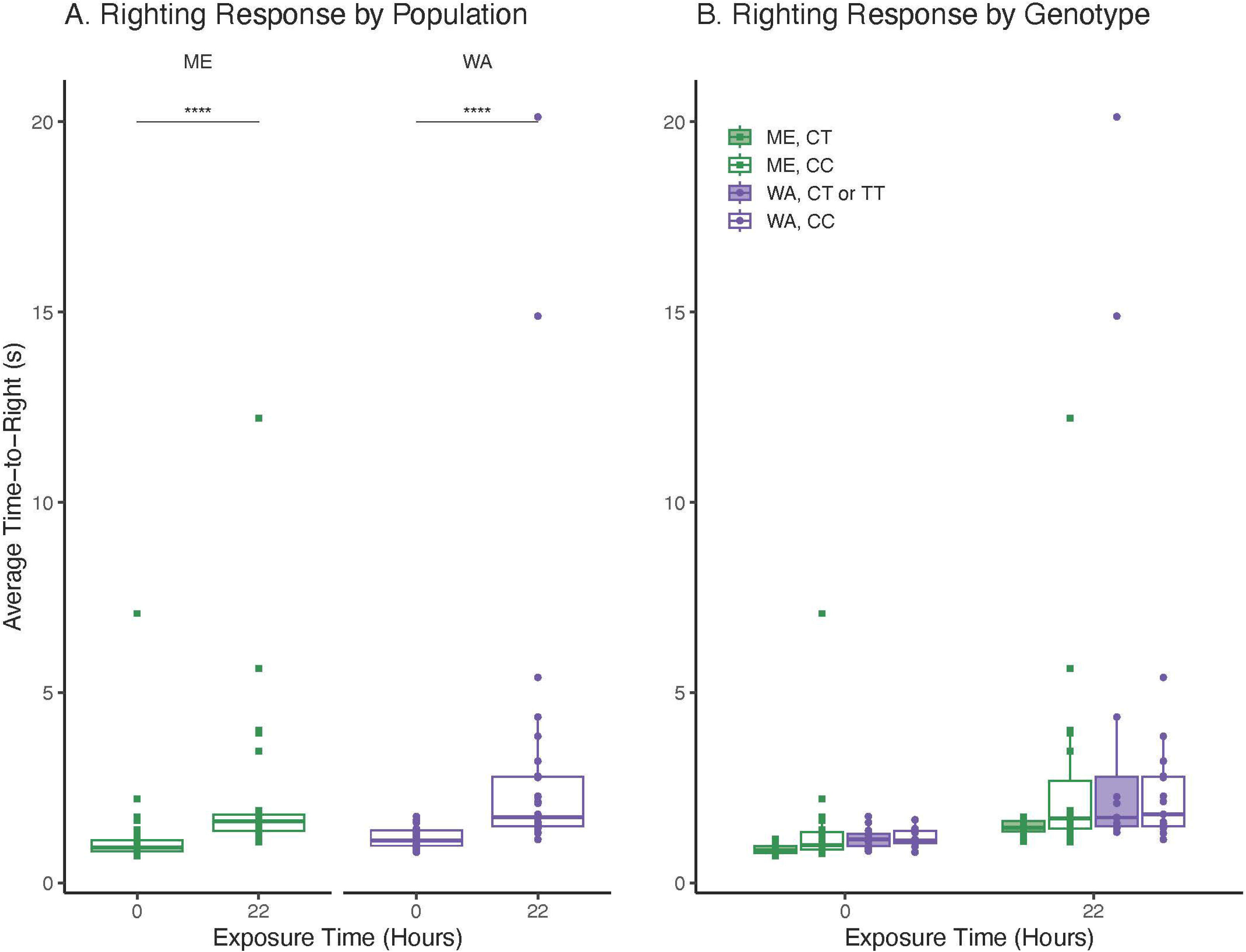
Righting response before (0 hours) and after (22 hours) heat shock by **A**) population and **B**) population and genotype. In all panels, TTR is represented by green squares for ME *C. maenas* and purple triangles for WA *C. maenas*. Significantly different pairwise comparisons are indicated on the figure (**** = *P*-value < 0.0001).

### Experiment 2: The role of “heat priming” in shaping thermal tolerance

The righting responses of *C. maenas* that previously experienced heat stress were compared to the responses of crabs naïve to stress in order to understand how repeated exposures impact thermal plasticity (**Figure S5**). We note that there were small temperature differences between tanks for the second heat shock (**Table S2**; **Figure S5**). Because of this experimental difference, tank was used as a random effect in subsequent analyses to account for significant temperature differences.

Average TTR for an individual crab was significantly impacted by treatment (X^2^ = 20.52, *P*-value = 0.0004), day (X^2^ = 24.16, *P*-value = 0.0008), and their interaction (X^2^ = 15.69, *P*-value = 0.004) (**Figure 3A**). These trends were driven by differences in TTR between control and treatment crabs at the end of the experiment. On day 4, *C. maenas* exposed to a prior heat shock had faster and less variable righting times (mean_treatment_ = 1.18 seconds ± 0.62 seconds) than their counterparts that did not experience a prior heat shock (mean_control_ = 4.78 s ± 4.98 s) (**Figure 3A**; t_control-treatment_ = 4.48, *P*-value_control-treatment_ = 0.001). Within-treatment comparisons revealed that crabs that did not experience a prior heat shock had significantly slower righting response on day 4 than they did on days 0 or 1 (**Figure 3A**; t_day0-day4_ = −4.02, *P*-value_day0-day4_ = 0.0005; t_day1-day4_ = −5.06, *P*-value_day1-day4_ < 0.0001).

**Figure 3.**
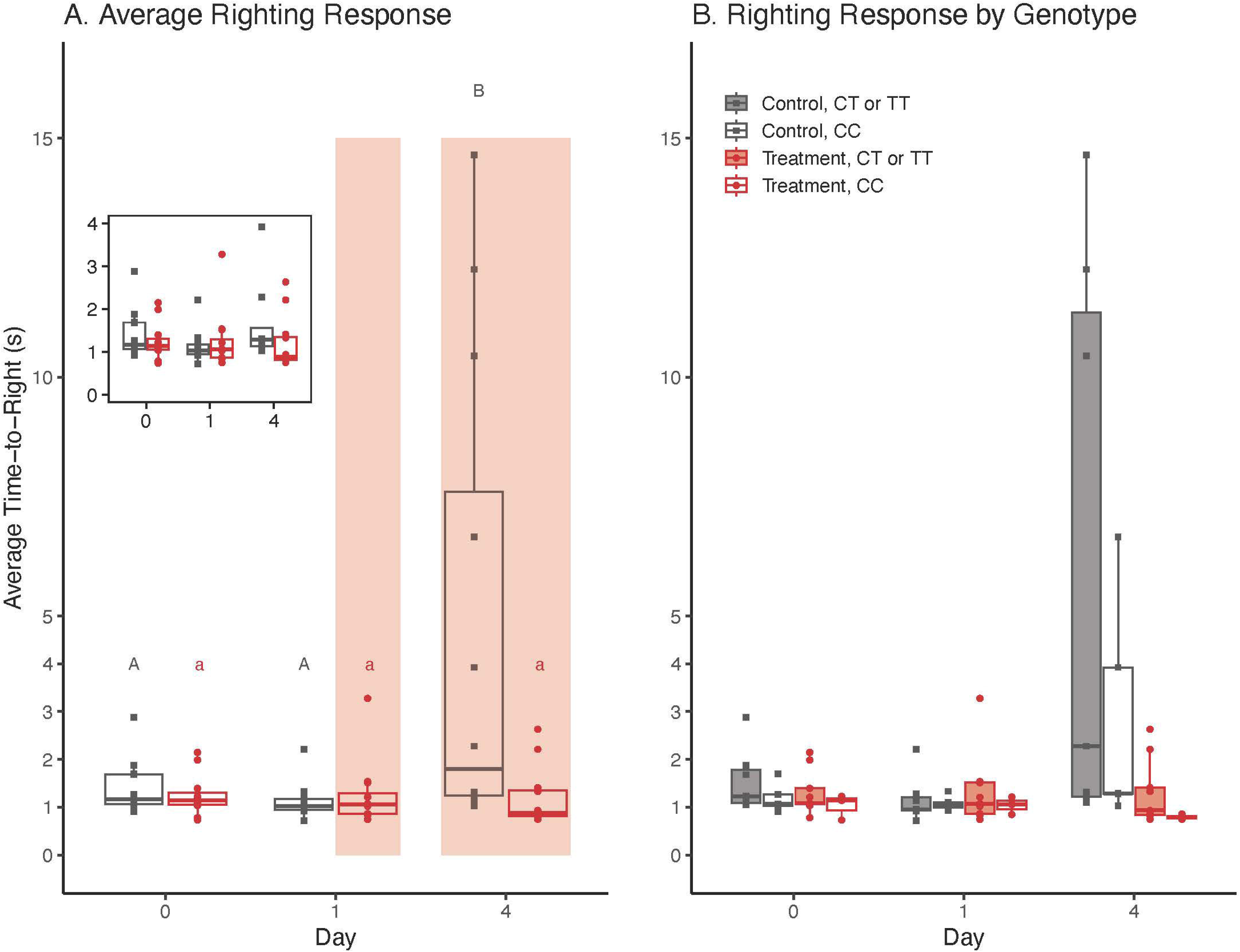
**A**) Average TTR across all timepoints for control (grey) and treatment (red) crabs. The inset shows data for crabs whose TTR was < 4 seconds. Timing of the heat shocks are indicated by transparent red boxes. Day 1 measurements were taken after the first heat shock for the treatment crabs, and day 4 measurements were taken after the second heat shock experienced by all crabs. Pairwise differences between days are indicated within control (grey uppercase) and treatment (red lowercase) treatments. **B**) Average TTR by genotype.

Supergene genotype was not associated with thermal response and priming. Genotype (X^2^ = 3.52, *P*-value = 1), presence of the C allele (X^2^ = 1.67, *P*-value = 1), and presence of the T allele (X^2^ = 2.73, *P*-value = 1) had no significant impact on average TTR. Although these trends were not statistically significant, crabs with at least one T allele had slightly slower TTR than their counterparts without a T allele, irrespective of prior heat shock exposure (**Figure 3B**; mean_treatment,day 4, T_ = 1.13 s ± 0.68 s, mean_treatment,day 4, no T_ = 0.80 s ± 0.06 s, mean_control, day 4, T_ = 6.17 s ± 6.01 s, mean_control, day 4, no T_ = 2.83 s ± 2.44 s). Crab ID and tank accounted for 23.41% or 0% of the variance in TTR in the final model (treatment, time, and their interaction only), respectively.

### Experiment 3: Changing cold tolerance across populations over time

Crabs from MA and WA were ramped down to target temperatures of 1.5°C or 5°C and held for several days to understand the factors that govern short- and medium-term cold tolerance (**Figure S7**). All tanks reached experimental temperatures by hour 22 of the 94 hour experiment. Actual temperatures experienced were slightly higher than the targets (**Table S2**). Differences in temperature conditions between these two experiments were likely due to the additional tanks used for WA and higher ambient temperatures in July 2024 versus June 2024 impacting cooling efficiency of the environmental chamber.

Righting response of MA crabs was significantly impacted by temperature (X^2^ = 77.69, *P*-value < 0.0001), time (X^2^ = 161.78, *P*-value < 0.0001), and their interaction (X^2^ = 51.81, *P*-value < 0.0001) (**Figure 4A**). Crabs at 1.5°C had significantly slower TTR than crabs in 5°C at all timepoints except for hour 0, when all crabs were at their common 15°C acclimation temperature (**Table S4**). Within each treatment, there were also significant differences in TTR between timepoints (**Figure 4A; Table S5**). Supergene genotype (X^2^ = 0.61, *P*-value = 1), presence of the C allele (X^2^ = 0.38, *P*-value = 1), or presence of the T allele (X^2^ = 0.19, *P*-value = 1) did not significantly impact average TTR in MA *C. maenas* (**Figure 4B**). Crab ID explained 10.99% of the variance in the final model (temperature, time, and their interaction only).

**Figure 4.**
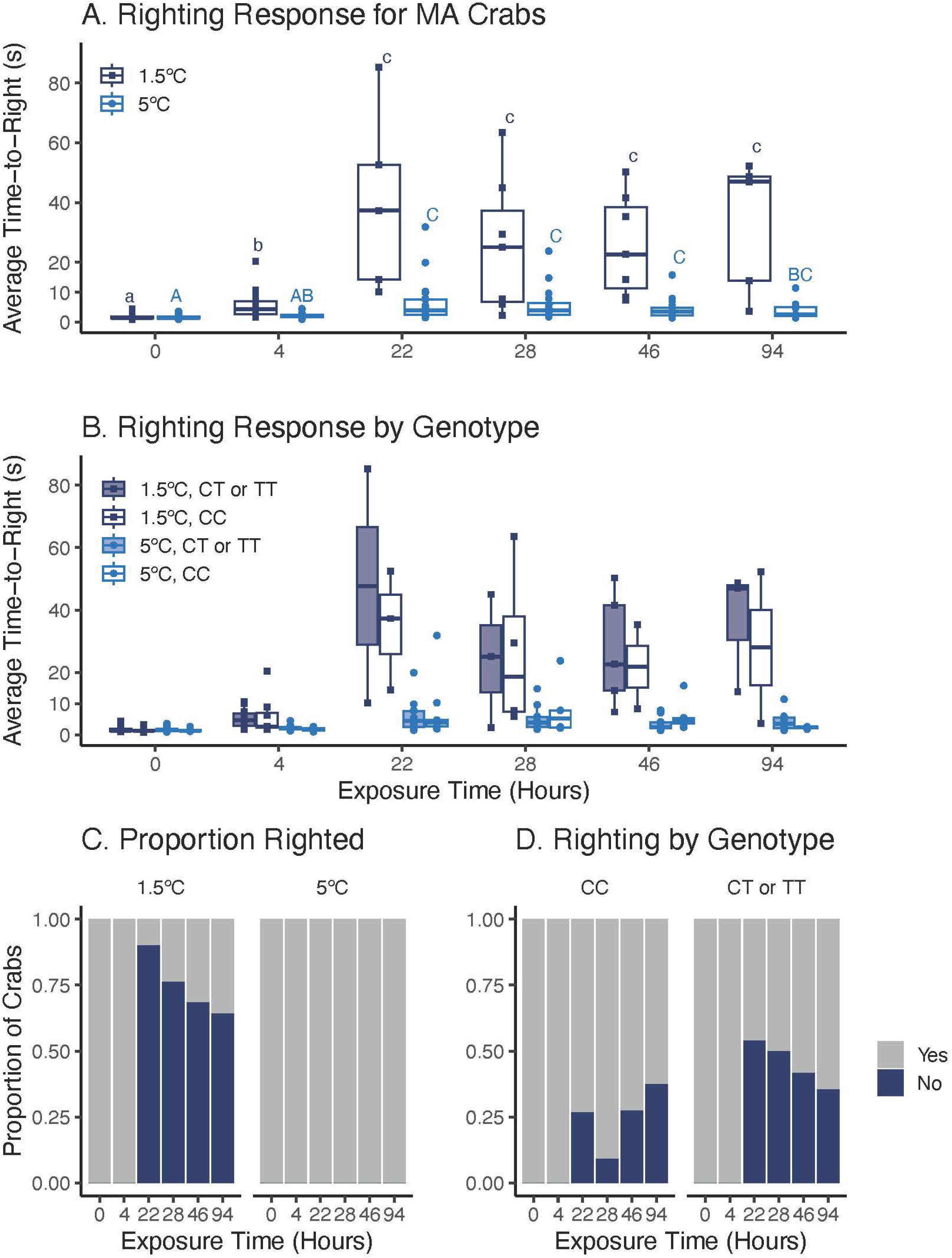
**A**) Average time-to-right for MA *C. maenas* in 1.5°C (dark blue squares) and 5°C (light blue circles) temperature treatments; crabs that failed to right in 90 s were not included in these data Righting response was significantly impacted by temperature, time, and their interaction. Pairwise differences between timepoints are indicated within 1.5°C (dark blue lowercase) and 5°C (light blue uppercase) treatments. **B**) Average TTR by genotype. **C**) Proportion of MA crabs that righted within 90 seconds. **D**) Proportion of MA crabs at 1.5°C that failed to right with and without a T allele. In panels **C**) and **D**), the dark blue color represents crabs that failed to right in 90 s. Failure to right was significantly impacted by temperature and time.

Throughout the experiment, some MA *C. maenas* failed to right at 1.5°C, but none failed to right at 5°C. Shortly after first reaching 1.5°C at hour 22, 90% of crabs failed to right, while 64.3% of the remaining crabs failed to right at the end of the experiment (**Figure 4C**). Failure to right was significantly associated with temperature (X^2^ = 56.05, *P*-value < 0.0001) and time (X^2^ = 64.72, *P*-value < 0.0001), but not their interaction (X^2^ = 1.07 × 10^-11^, *P*-value = 1), genotype (X^2^ = 3.31, *P*-value = 1), nor the presence of the C allele (χ^2^ =1.99 × 10^-11^, *P*-value = 1). The majority of crabs that failed to right had at least one T allele (77.8% of crabs at 22 hours, 66.7% of crabs remaining at 94 hours) (**Figure 4D**). However, the impact of the T allele on failure to right in the MA population was not significant after multiple test correction (X^2^ = 3.31, *P*-value = 0.82). Failure to right was not impacted by sex, integument color, carapace width, weight, or whether or not a crab was missing a fifth pereopod in any population (**Table S7**). There were no differences between the final model and the most parsimonious model derived from the sensitivity test for either MA model (**Table S8**).

Crabs from WA responded to 1.5°C and 5°C in a similar fashion to their MA counterparts. Temperature (X^2^ = 113.80, *P*-value < 0.0001), time (X^2^ = 287.84, *P*-value < 0.0001), and their interaction (X^2^ = 65.95, *P*-value < 0.0001) significantly impacted average righting response in WA *C. maenas* (**Figure 5A**). Righting response in 1.5°C was significantly slower than TTR in 5°C for all timepoints except hour 0, prior to the start of the cold ramp (**Table S5**). Within a treatment, there were also significant differences in righting response over time (**Table S6**). There was no significant impact of genotype (X^2^ = 1.06, *P*-value = 1), presence of the C allele (X^2^ = 0.40, *P*-value = 1), or presence of the T allele (X^2^ = 0.54, *P*-value = 1) on average TTR in WA crabs (**Figure 5B**). Righting response was not impacted by sex, integument color, carapace width, weight, or whether or not a crab was missing a fifth pereopod (**Table S3**). Crab ID explained 5.32% of variance in the final model (temperature, time, and their interaction only). There were no differences between the final model and the most parsimonious model derived from the sensitivity test (**Table S6**).

**Figure 5.**
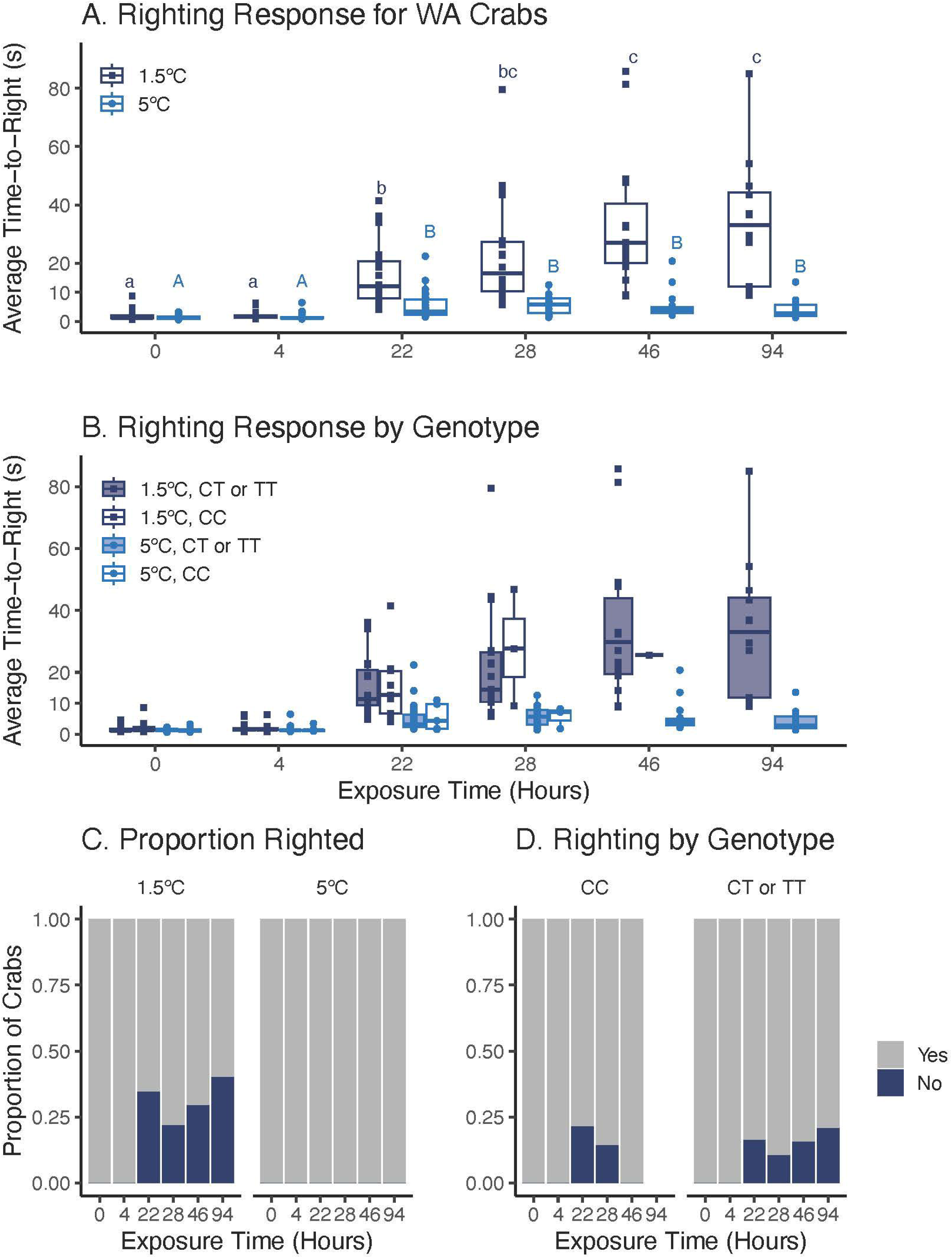
**A**) Average time-to-right for WA *C. maenas* in 1.5°C (dark blue squares) and 5°C (light blue circles) temperature treatments; crabs that failed to right in 90 s were not included in these data. Righting response was significantly impacted by temperature, timepoint, and their interaction. Pairwise differences between timepoints are indicated within 1.5°C (dark blue lowercase) and 5°C (light blue uppercase) treatments. **B**) Average TTR by genotype. **C**) Proportion of MA crabs that righted within 90 seconds. **D**) Proportion of MA crabs at 1.5°C that failed to right with and without a T allele. In panels **C**) and **D**), the dark blue color represents crabs that failed to right in 90 s. Failure to right was significantly impacted by temperature and time.

In comparison to the MA population, a smaller proportion of WA *C. maenas* failed to right at 1.5°C: 34.5% WA crabs failed to right at hour 22 and 40% of the remaining crabs failed to right at hour 94 (**Figure 5C**). Failure to right in WA crabs was related to temperature (X^2^ = 27.10, *P*-value < 0.0001) and time (X^2^ = 23.46, *P*-value < 0.0001), but not their interaction (X^2^ = 0, *P*-value = 1). There was also no impact of genotype (X^2^ = 2.64, *P*-value = 1), presence of the C allele (X^2^ = 2.60, *P*-value = 1), or presence of the T allele (X^2^ = 0.06, *P*-value = 1) on failure to right in WA *C. maenas*. Time did not have a significant impact in the most parsimonious model derived for the sensitivity test, and was therefore removed from the final model for the full dataset (**Table S8**). Temperature was still significant in the revised model (X^2^ = 26.57, *P*-value < 0.0001).

Population (MA or WA) did not have a significant impact on average TTR (X^2^ = 0.47, *P*-value = 0.49) or overall failure to right (X^2^ = 0, *P*-value = 1). Population-specific impacts on failure to right at 1.5°C were also examined for crabs at each timepoint. There was no significant impact of population at any timepoint (*P*-value > 0.05 for all comparisons).

## Discussion

We examined tolerance and phenotypic plasticity in response to heat and cold exposures in a highly-successful species in two regions where it has been introduced and lost significant genetic diversity relative to its native range. We found that acute exposure to sub-lethal high temperature elicits a stress response which could be ameliorated by repeated sub-lethal stress exposure. Exposure to cold temperatures led to slower righting or failed righting. However, some crabs were able to recover, suggesting longer-term plasticity at cold temperatures. Interestingly, a genomic region strongly associated with temperature and thermal physiology at the population level showed no relationship with thermal physiology at the individual level in our experiments. The maintenance of broad thermal tolerance and plasticity in righting response in adult crabs with no apparent influence of supergene genotype suggests that adult tolerance is largely reliant on outcomes of selection during early life history in addition to robust phenotypic plasticity.

### Green crabs display adaptive plasticity at sub-lethal temperatures

Our work investigating *C. maenas* response to heat shock adds to the body of work highlighting the broad thermal tolerance of this species outside of its native range. We expected that *C. maenas* would speed up righting response during the heat shock because metabolic rates increase for ectotherms at warmer temperatures (Somero, Lockwood and Tomanek 2017). Instead, we found that crabs from ME and WA slowed TTR after heat shock. This contrasts with Jost et al. (2012), who report TTR starting to slow at 34°C for ME *C. maenas*, with no decrease at 32°C. Methodological differences may be responsible for these contrasting results. While our study held crabs at 30°C for ∼22 hours after a ∼1°C/hour heat ramp, Jost et al. (2012) used a 6°C/hour increase in temperature. The faster temperature used in Jost et al. (2012) may activate different acute response pathways, leading to a different physiological outcome. Alternatively, the slower TTR observed in our study could indicate that thermal thresholds may be reduced with prolonged exposure to an elevated temperature. Interestingly, crabs in the treatment group experiencing their first 30°C heat stress on day 1 in Experiment 2 did not exhibit stress-related slowing of righting response, while crabs in the control group experiencing a heat shock on day 4 did significantly slow their righting response. This unexpected result could be the product of individual variation in stress response within our sampled population: crab ID contributed to ∼20% of variance in righting response, which was the highest out of any experiment. Future work should repeat this experiment with a larger sample size and a range of heat shock timing and duration to clarify the impact of initial heat stress on physiology.

The response of WA crabs to repeated heat stress demonstrates that negative impacts of acute sub-lethal stress may be ameliorated by adaptive plasticity. This adaptive plasticity may manifest in different ways, such as production of molecular chaperones like heat shock proteins (HSPs) and transcription factors (Somero, Lockwood and Tomanek 2017). After the heat shock subsides, these heat shock-responsive molecules and regulatory pathways may remain active in the organism, preparing it for similar conditions within a certain timeframe. While not widely explored in crustaceans, beneficial priming (*ex*. heat hardening or hormetic priming) has been documented in arthropods such as *Drosophila melanogaster* (Sejerkilde, Sørensen and Loeschcke 2003; Sørensen *et al*. 2008; Moghadam *et al*. 2019), as well as a variety of marine taxa, including seagrasses (Pazzaglia *et al*. 2021), anemones (Glass *et al*. 2023), corals (Martell 2022), shellfish (Abbas *et al*. 2024), and sticklebacks (Spence-Jones *et al*. 2025). Studies have also shown that priming may be a contributing factor to increased thermal tolerance in non-indigenous species. Repeated heat stress increased thermal tolerance in *Semimytilus algosus* and *Mytilus edulis* mussels (Lenz *et al*. 2018) and priming improved heat tolerance in non-native *Bactrocera dorsalis* and *Bactrocera correcta* flies (Gu *et al*. 2019). The conditions under which beneficial plasticity manifests should be evaluated in future experiments by altering the intensity of heat shocks as well as the time in between the first and second heat shocks.

### Righting response is plastic after short- and medium-term cold exposure

Albeit less frequently studied than heat tolerance, cold tolerance often plays an important role in dictating species distributions (Wasson *et al*. 2025). Organisms that are able to tolerate cooler conditions may be able to effectively colonize northern latitudes with limited competition (Walther *et al*. 2009; Christiansen *et al*. 2015; Dijkstra, Westerman and Harris 2017). Our work demonstrates adult green crabs can maintain motor function at 5°C — with some maintaining motor function at 1.5°C — which may allow them a competitive advantage in colder conditions. These findings are in line with previous research that highlights the extensive cold tolerance of adult *C. maenas*. For example, *C. maenas* successfully reproduce in sub-arctic conditions in Newfoundland (Best, McKenzie and Couturier 2017), and can survive at 2°C for at least two months (Rivers, McKenzie and McGaw 2025), and below 5°C for over four months (Kelley, de Rivera and Buckley 2013). In prior work in a UK population in the native range, some crabs failed to right at 0°C, while crabs that maintained movement had a ∼5 second average righting response at 0°C (Young, Peck and Matheson 2006). We observed an average righting response of 33.09 s ± 22.58 s (MA) or 33.59 s ± 22.29 s (WA) at 1.5°C. These apparent differences in cold tolerance between crab populations may be due in part to genetic or environmental differences. However, it is likely that methodology plays a role. We included hesitation time, or the time before moving the fifth pereopod pair after crabs are placed dorsal side up, as part of the righting response, while Young et al. (2006) does not, which likely increased our measurements relative to theirs. Nevertheless, it is clear that *C. maenas* has a robust cold tolerance, which may have facilitated northern range expansion into Newfoundland (Blakeslee *et al*. 2010) and ongoing spread in Alaska (NOAA 2022).

In addition, *C. maenas* displayed some ability to acclimate to chronic cold stress. At 5°C, righting response for several MA and WA crabs sped up at the end of the experiment relative to earlier timepoints (**Figure S8**). We also observed evidence of acclimation to 1.5°C: while many crabs failed to right when initially exposed to 1.5°C, some individuals “recovered” and were able to right at a later timepoint (**Figure S8**). This recovery suggests that failure to right is thermally plastic, and may be related to dormancy in this species. Historical work suggests that when temperature is ≤5°C *C. maenas* enter torpor (Berrill 1982; Young, Peck and Matheson 2006), an energy conservation mechanism typically employed in colder conditions (Pörtner and Playle 1998). Recent work suggests that crabs enter a dormant state (metabolic slowing with limited movement) which may facilitate energy conservation while also allowing for opportunistic feeding and locomotive activity (*ex*. burying for predator avoidance) (Rivers, McKenzie and McGaw 2025). The use of dormancy is also supported by prior research which found a steady decrease in heart rate with no cardiac breakpoint when cooling crabs from 15°C to 0°C, suggesting that *C. maenas* has sub-zero cold tolerance (Tepolt and Somero 2014). Adult locomotive plasticity in cold temperatures may be facilitated by additional mechanisms such as regulation of hemolymph magnesium to prevent calcium channel blockage and restore locomotor activity (Frederich *et al*. 2000; Aronson *et al*. 2015); increasing the number of polyunsaturated lipids present in muscle tissue to provide membrane fluidity (Chapelle 1978); and downregulating the cell cycle regulator *cyclin D1* to reduce cell proliferation (Kelley, de Rivera and Buckley 2013). In addition to exploring the mechanisms that govern adult dormancy in cold conditions, attention should be paid to cold tolerance mechanisms throughout life history. Larval development and survival in *C. maenas* are compromised when temperatures are below 10°C (Nagaraj 1993; Hines *et al*. 2004; deRivera *et al*. 2007), and modeling work suggests that cold temperatures have played a key in limiting larval survival and spread in the northeast Pacific (Du *et al*. 2024). Additional manipulative experiments with larval crabs may clarify differences in cold tolerance associated with ontogeny.

We explored the relationship between supergene and cold acclimation at the individual level in part because of a significant population-level association between supergene allele and cold tolerance in prior work (Tepolt and Palumbi 2020). Surprisingly, in our study the cold-adapted supergene did not significantly impact TTR at 1.5°C or 5°C. This is consistent with Coyle et al. (2019), who found that failure to right after acute exposure to 4.5°C was associated with mitochondrial haplotype, not supergene genotype, in male crabs from Gulf of Maine populations. Even though supergene distributions were similar between MA and WA populations, the MA population had a higher proportion of crabs that failed to right during the experiment. While these differences among populations may be a product of the slightly different experimental temperatures for MA and WA crabs, it is unlikely that this ∼0.5°C temperature difference is solely responsible. The switch from an active to dormant state in *C. maenas* occurs between 4-6°C (Young, Peck and Matheson 2006; Rivers, McKenzie and McGaw 2025), which is well above the coldest temperature used in this experiment. While mitochondrial diversity present in the northwest Atlantic could interact with nuclear diversity to shape thermal tolerance, to our knowledge there is limited mitochondrial haplotype diversity within the MA population (Darling *et al*. 2014; Jeffery *et al*. 2017; Lehnert *et al*. 2018). Instead, we suggest that MA crabs utilize physiological mechanisms that elicit a more pronounced dormancy than WA crabs as a response to colder winters in the northwest Atlantic (**Figure 1**). Even though we acclimated all crabs to the same conditions prior to experimentation, there is evidence that distal environmental history can impact physiological response through within- and across-generation legacy effects in marine invertebrates (Donelan *et al*. 2020; Ashey and Rivest 2021; Fernández *et al*. 2025). Future work should account for legacy effects, and examine responses of crabs during the winter season to ascertain if population differences are a result of environmental history or lack of seasonal acclimation.

## Conclusion

Our work highlights how thermal plasticity contributes to the broad thermal tolerance of successful non-indigenous species, even after a substantial loss of genetic diversity. We found that crabs exhibit potentially beneficial plasticity in response to sub-lethal heat stress. Additionally, crabs display a robust cold tolerance and the potential to acclimate to cold temperatures over the course of 94 hours. Although previous work demonstrates a strong association between supergene allele frequency and both cold tolerance and sea surface temperature at the population level (Tepolt and Palumbi 2020) This suggests that, rather than being causally linked, environmental conditions like temperature likely shape both supergene distribution and phenotypic plasticity throughout life history, including during larval dispersal and settlement. In order to clarify the relationship between population- and individual-level tolerance, future work should compare if and how the supergene influences thermal tolerance in planktonic larvae, newly-settled juveniles, and adult crabs.

While we didn’t see significant relationships between genotype and individual righting response, it is possible that supergene genotype plays a role at a different level of biological organization. Since the association was established using heart rate at 0°C (Tepolt and Palumbi 2020), it may be useful to examine the relationship between genotype and heart rate plasticity in individual crabs. Supergene genotype could also impact molecular pathways without changes in whole-organism physiology. For example, a study in coral larvae detected metabolic reprogramming under elevated temperature without a decrease in survival (Huffmyer *et al*. 2024), and American lobster (*Homarus americanus*) exposure to ocean acidification resulted in broad metabolic reprogramming that was not associated with changes to resting metabolism (Noisette *et al*. 2021). Future work should examine if the warm- and cold-adapted supergene alleles are associated with differences in metabolic pathways and gene expression that allow for physiological homeostasis.

## Author contributions

YRV, JCK, and CP conceived, designed, and carried out all experiments with assistance from HLF and input from CKT. YRV, JCK, and JK conducted molecular assays. JCK, CP, and HLF conducted preliminary data analysis. YRV refined all data analysis. YRV wrote the initial draft with input from JCK and CP; and YRV and CKT revised and edited this manuscript. All authors reviewed and approved the final manuscript.

## Acknowledgements

We are grateful to Chelsey Buffington and Brian Turner at Washington Fish and Wildlife and Tait Nygaard and Isabelle See at Quahog Bay Conservancy for providing crabs used in this experiment, and the PEP and CC-CREW interns that volunteered with data collection. We also appreciate two anonymous reviewers for their constructive feedback on an earlier draft.

## Data availability

All raw data, scripts, results, and supplementary material are available at https://github.com/yaaminiv/cold-green-crab.

