## Supplementary figures and images for "Plasticity, not genetics, shapes individual responses to thermal stress in non-native populations of the European green crab (*Carcinus maenas*)"

### Figure S1

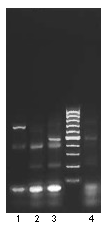

### Figure S2

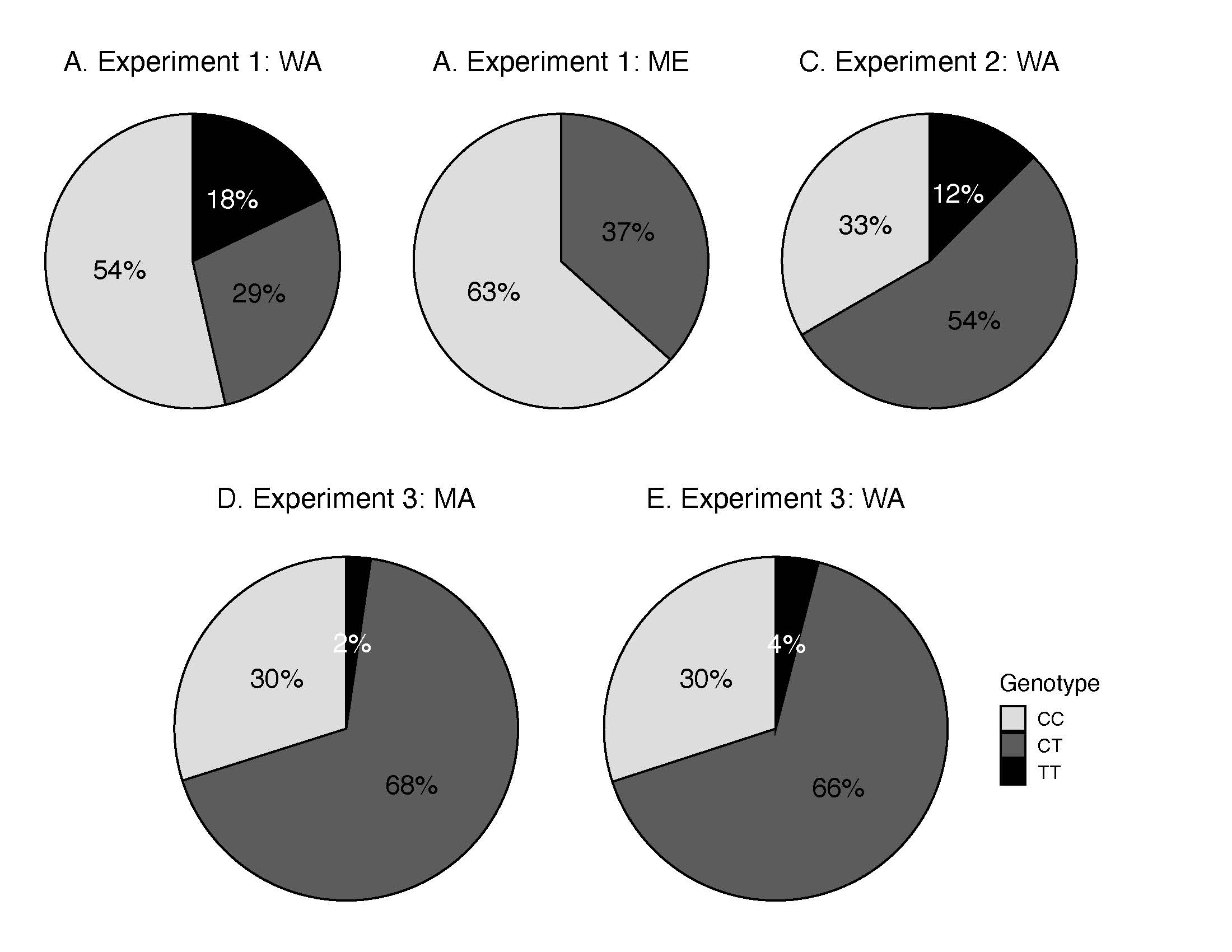

### Figure S3

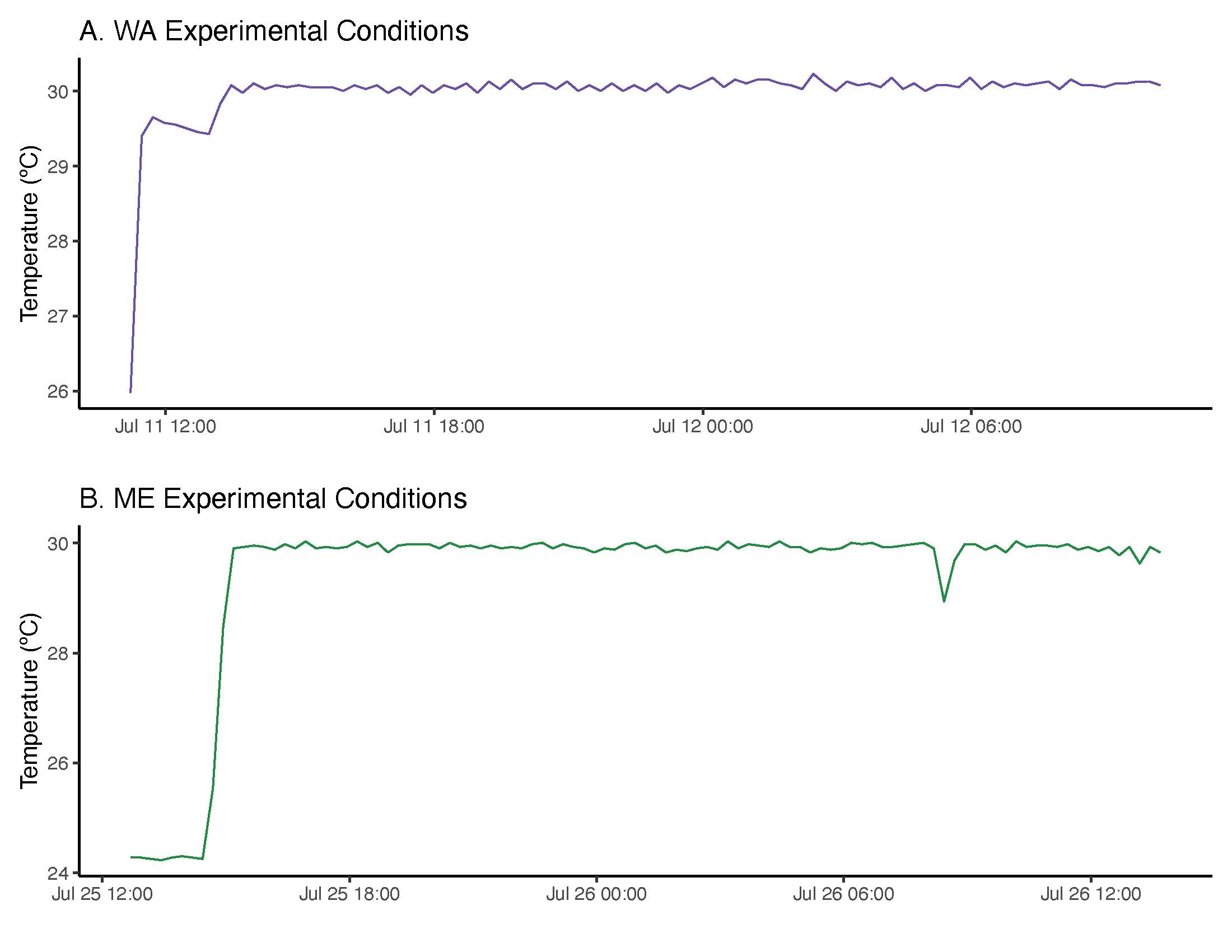

### Figure S4

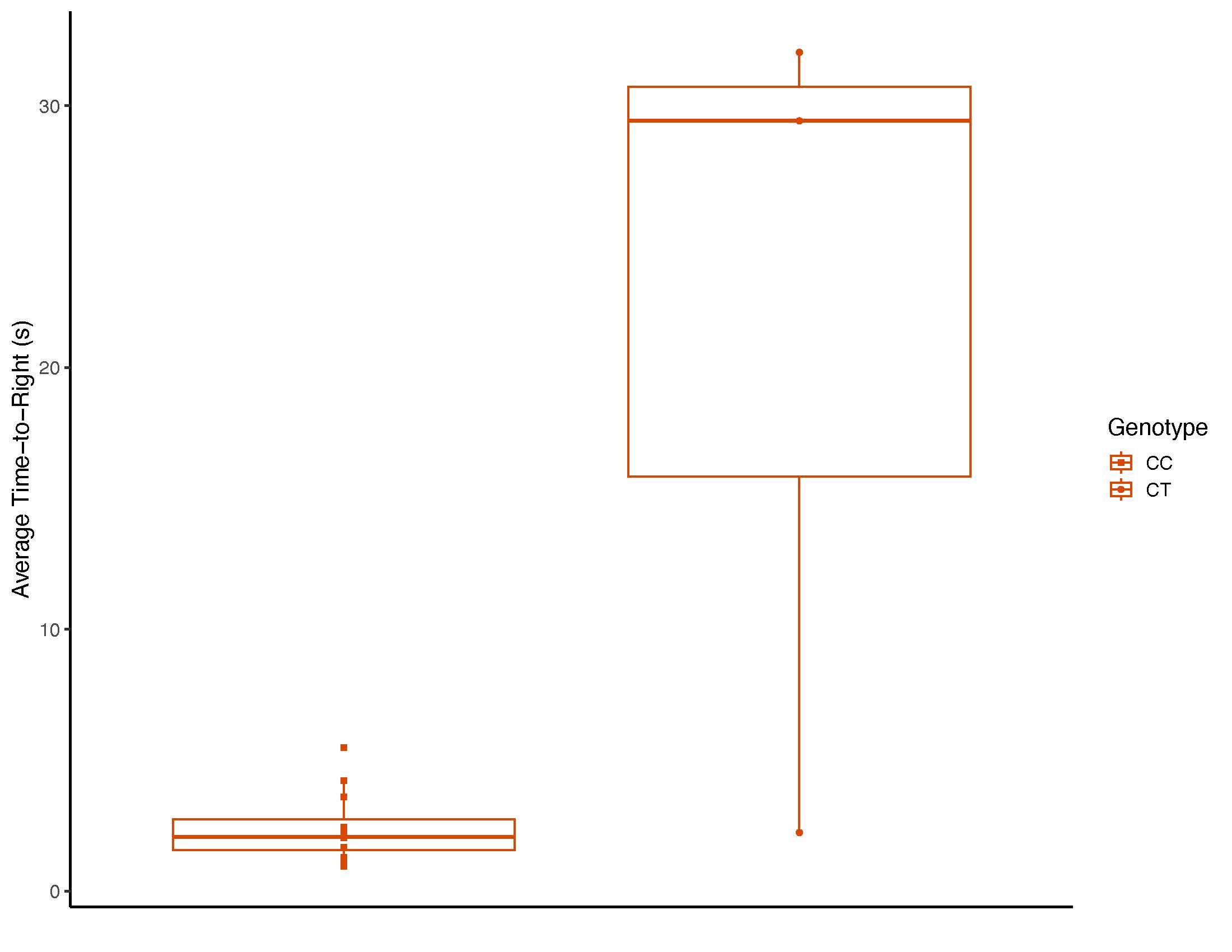

### Figure S5

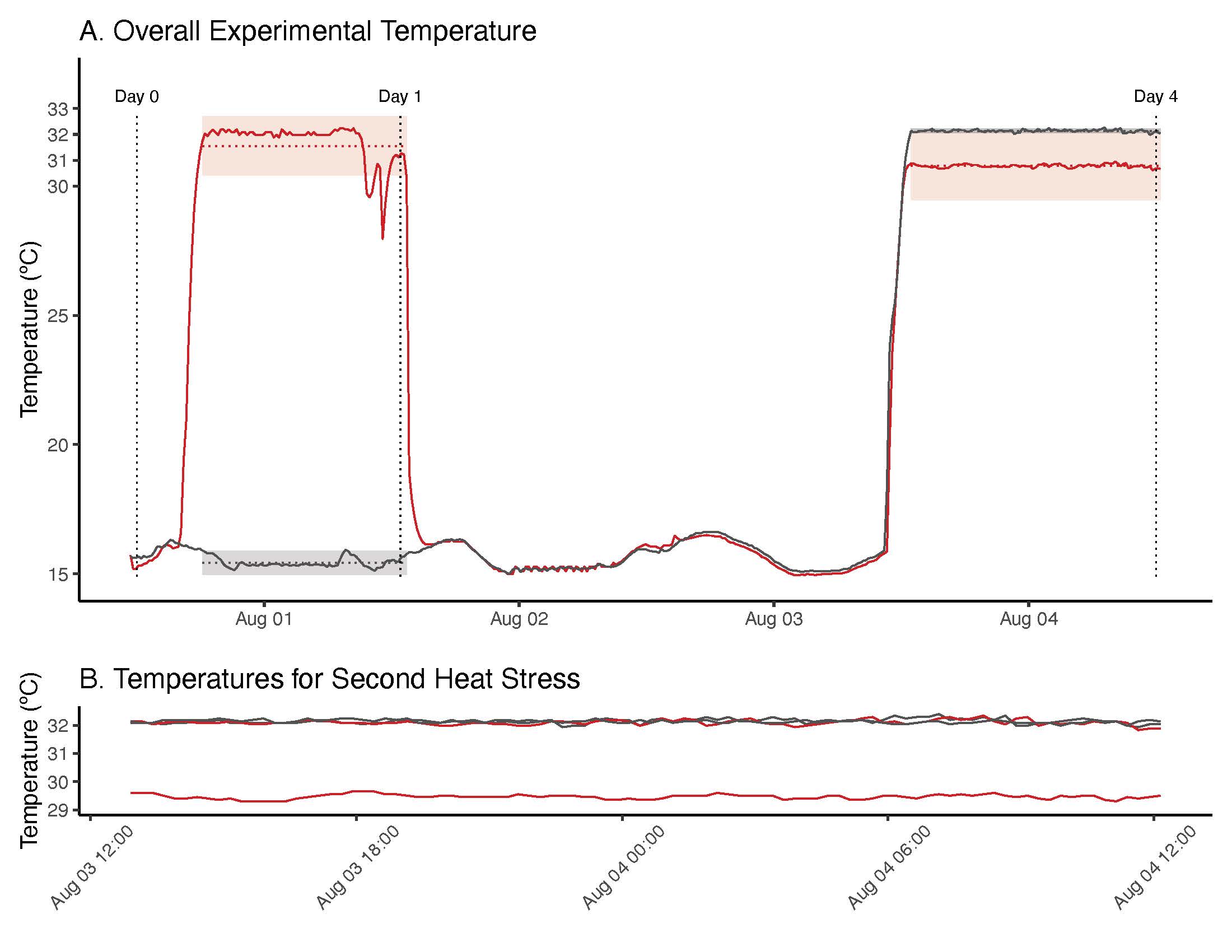

### Figure S6

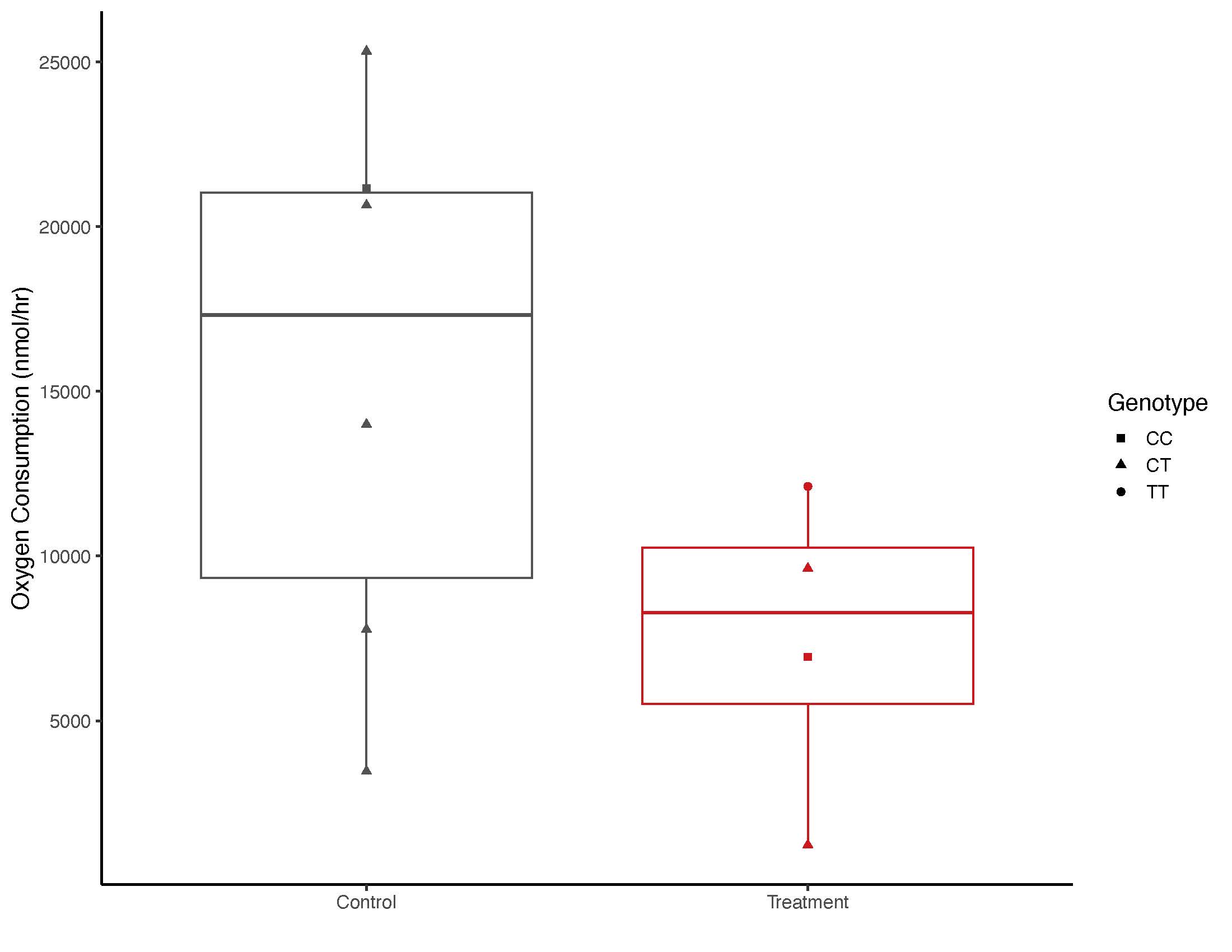

### Figure S7

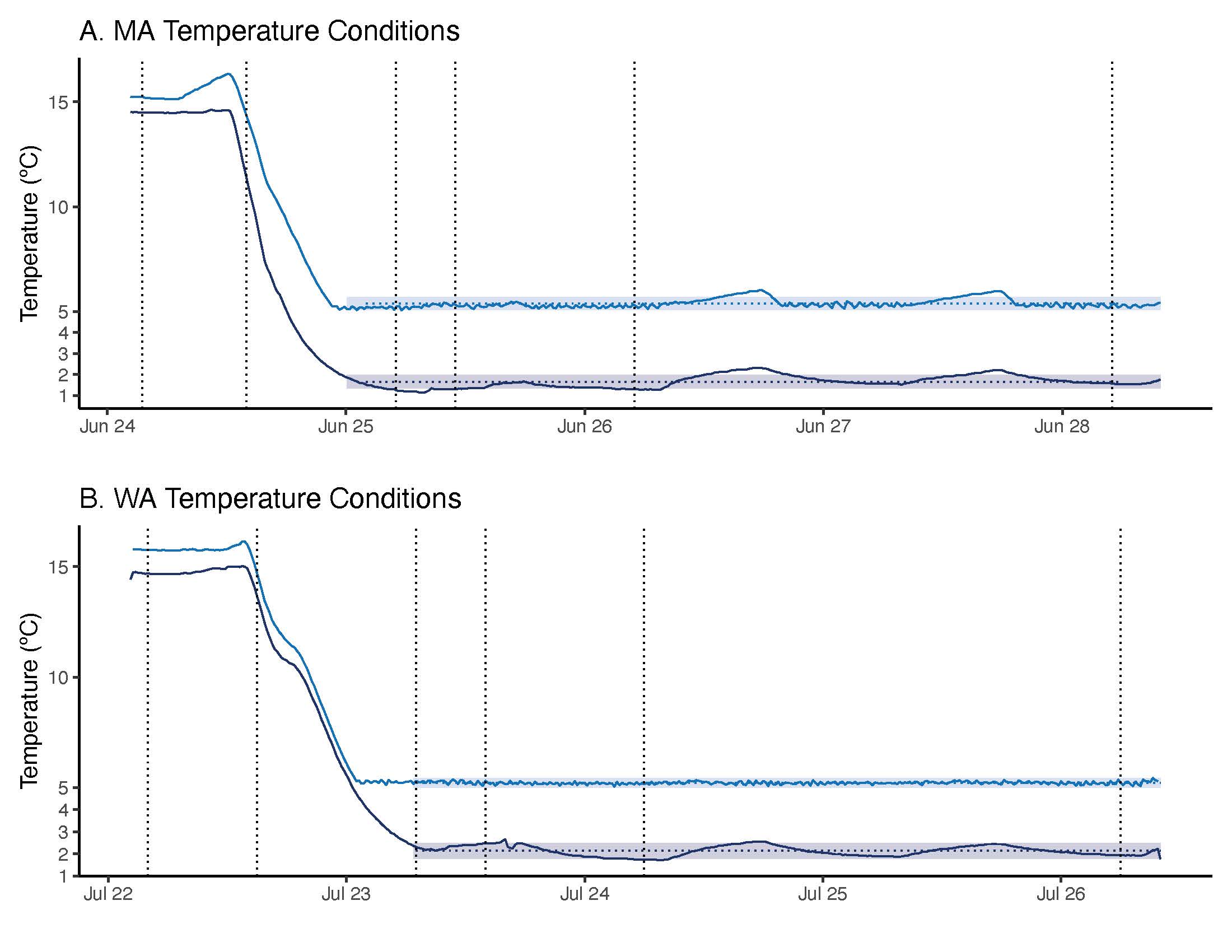

### Figure S8

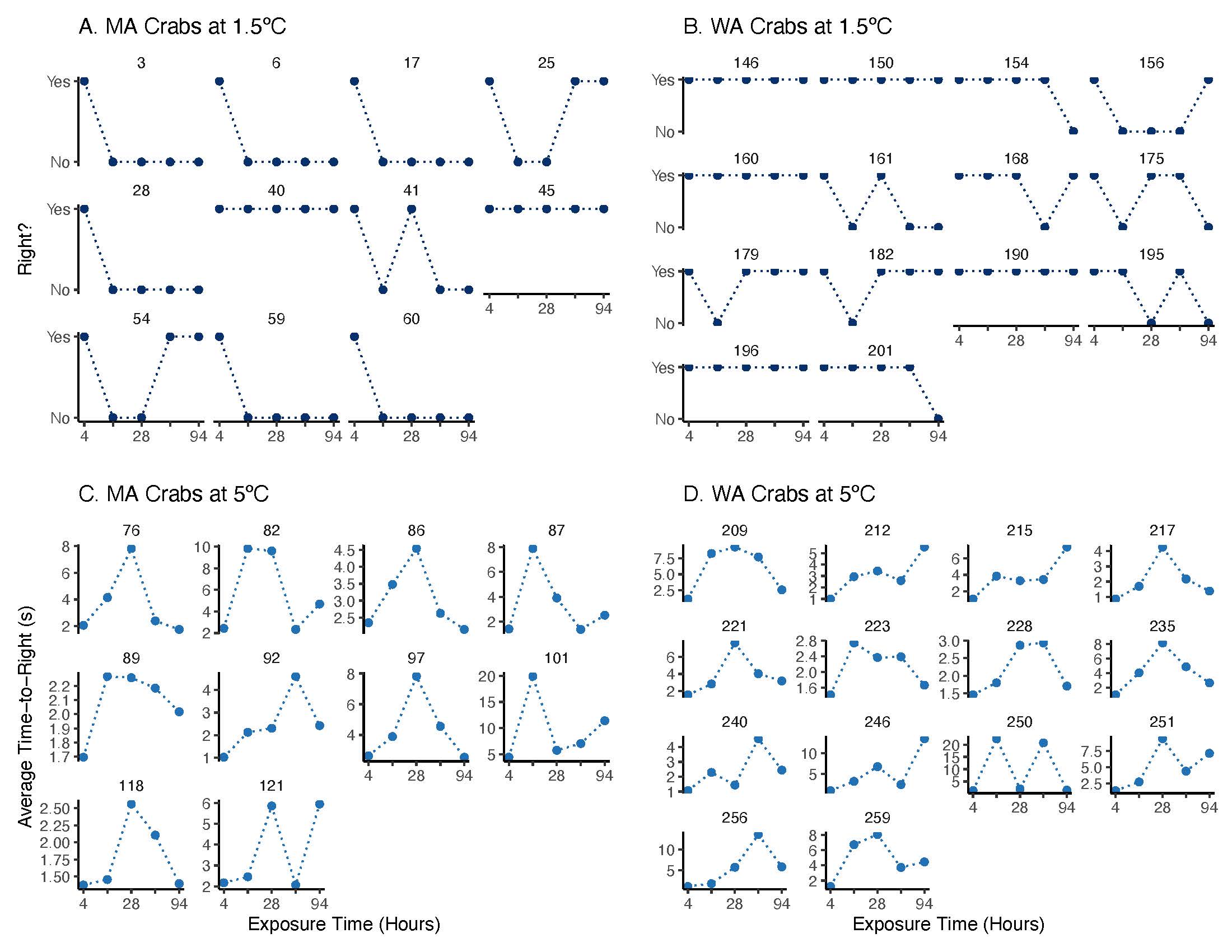
