## Appendix for "Plasticity, not genetics, shapes individual responses to thermal stress in non-native populations of the European green crab (*Carcinus maenas*)"

### Supplementary Information

#### Appendix S1: Green crab invasion history and genetic diversity in North America

Green crabs experienced an initial genetic bottleneck upon establishment in the northwest Atlantic in the early 1800s (Carlton and Cohen 2003; Roman and Palumbi 2004; Roman 2006; Darling *et al.* 2008; Tepolt and Palumbi 2015), with populations in the northeast Pacific experiencing a second bottleneck after being introduced from a northwest Atlantic source (Carlton and Cohen 2003; Tepolt *et al.* 2009; Tepolt and Palumbi 2015). A second introduction of crabs from the northern part of the native range may have introduced genetic variation associated with more robust cold tolerance, allowing the crabs to expand into Maritime Canada and Newfoundland (Roman and Palumbi 2004; Roman 2006; Blakeslee *et al.* 2010; Weihrauch and McGaw 2024). However, the genetic legacy of this second introduction has largely been restricted to the northern extent of the species’ northwest Atlantic range. Despite this substantial loss of genetic variation across much of their range, crabs from several introduced North American populations maintained extensive plasticity of both heat and cold tolerance, comparable to that of crabs in the native range (Tepolt and Somero 2014a).

#### Appendix S2: General experimental setup

A series of experiments was conducted to assess the impact of short- (~24 hours) to medium-term (94 hours) heat and cold stress on introduced *C. maenas* populations in both the northwest Atlantic and northeast Pacific. Experimental crabs were individually labeled using waterproof paper and superglue, and the propus and dactyl segments from the third walking leg on the right side (dorsal view) were preserved in 95% ethanol for subsequent genotyping. These joints were taken as the third walking leg has no significant role in crab righting behavior (Young, Peck and Matheson 2006). If the crab did not have a third walking leg on the right side, the third walking leg was taken from the left.

Crabs were placed in 21 cm x 26 cm x 41 cm (22.386 L) tanks filled with 800 micron filtered raw seawater from Great Harbor, MA, USA (Experiments 1 and 3) or 50 micron filtered raw seawater from Vineyard Sound, MA, USA (Experiment 2). Each tank had an air stone (Quickun Aquarium Air Stone) attached to a 10L aerator (Whisper® Aquarium Air Pumps, Tetra®) and an aquarium filter (MARINA i25). Crabs were distributed to balance sex, carapace width, and integument color among tanks. Crabs with orange-red or red integument color were not included in experiments; these colors indicate an extended time in intermolt and “red” green crabs have less robust tolerance to a range of physiological stressors (Styrishave, Rewitz and Andersen 2004). Remaining crabs had integument colors at the beginning of the experiment ranging from blue to yellow-orange.

Tanks were checked for mortalities and fed daily. Throughout the acclimation and experimental periods every tank was given ½ teaspoon of commercial crab food (Crab Cuisine, Hikari USA Inc.) if tank temperature was ≥ 15ºC, and ¼ teaspoon of food if tank temperature was ≤ 5ºC. These quantities were sufficient to satiate crabs since there was always a small amount of uneaten food the next day. Particulate waste was removed prior to feeding. Every third day, ammonia concentrations were tested using an API® Ammonia Test Kit prior to removing particulate waste from tanks and performing a 60% water change. A conditioner (½ tsp Kordon Amquel Plus Aquarium Water Conditioner) was added to tanks during water changes if the ammonia concentration was above 1.0 ppm. Filter cartridges were replaced every five days A 12-hour:12-hour light:dark cycle was maintained for all tanks (Tepolt and Somero 2014b).

Crabs were initially kept in acclimation conditions for at least five days (see details below). Water temperature was maintained during the acclimation period using either a flowing water bath or environmental chamber manipulation. Water temperature in each tank was recorded every 15 minutes using two HOBO Data Loggers (Onset, USA), one at the water surface and one at the bottom of the tank. Warming (Experiments 1, 2, and 3) was performed using 300-Watt Deluxe Titanium Heating Tubes (Finnex, USA) with an external 1650-Watt temperature controller (bayite, China). Cooling (experiment 3) was performed in an environmental chamber with 0ºC as a set point. A non-parametric Kruskal-Wallis test (kruskal.test from R stats package (R Core Team 2024) was used to confirm that temperature treatments were statistically different from each other and that replicate tanks were not different from each other.

#### Appendix S3: Experiment 1: Responses to acute heat shock across populations

##### Methods

Crabs were sourced from Grays Harbor, WA, USA by Washington Fish and Wildlife on June 26, 2024 (n_WA_ = 30; carapace width_WA_ = 49 mm ± 6 mm, weight_WA_ = 25.11 g ± 11.35 g), or trapped at Harpswell, ME, USA by Quahog Bay Conservancy on July 18, 2024 under Maine Department of Marine Resources Green Crab Landing Licenses 11915 and 36329 (n_ME_ = 33; carapace width_ME_ = 51 mm ± 4 mm, weight_ME_ = 28.09 g ± 7.44 g) (**Figure S2A-B**). Experiments took place on July 11-12, 2024 for WA crabs and July 25-26 for ME crabs.

##### Results

Crabs from both WA and ME were exposed to a 22 hour heat shock to understand short-term responses to high temperatures. WA crabs were exposed to 30.04ºC ± 0.30ºC (Figure 2A), and ME crabs were exposed to 29.91ºC ± 0.30ºC (**Figure S3**; **Table S2**). No WA crabs died during the acclimation period or experiment, while four ME crabs (three CT, one TT) died during the acclimation period. Parasite intensity was quantified in WA and ME crabs to control for external factors impacting righting response and between-population comparisons. The hepatopancreas of ME *C. maenas* contained 0-105 metacercarial cysts (intensity = 0.46 cysts/mm ± 0.56 cysts/mm), while were not found in the hepatopancreas tissue of crabs from WA. No other parasites were identified in these populations.

##### Genotype impacts on righting response in New Hampshire population

The influence of supergene genotype on righting response in room temperature conditions was evaluated using *C. maenas* trapped at New Castle, NH, USA on July 22-24, 2024 under NH permit MFD 2426 (n_NH_ = 15; carapace width_NH_ = 69 mm ± 7 mm, weight_NH_ = 73.25 g ± 21.16 g). Crabs were acclimated at room temperature (~22ºC) for five days before experimentation. A generalized linear model was used to assess the impact of genotype on log-transformed average righting response, with *P*-values adjusted using the Bonferroni correction.

Crabs were dissected per approach in Experiment 1 to characterize the macroparasite community in this population. Crabs from NH were parasitized by either trematode metacercaria, acanthocephalans, or both. Individual crabs contained between 0-471 trematode metacercarial cysts and 0-19 acanthocephalans. Parasite intensities were 1.06 ± 2.20 cysts/mm and 0.04 ± 0.07 parasites/mm for trematode metacercaria and acanthocephalans, respectively. Neither trematode metacercarial cyst infection intensity (t = 1.42, *P*-value = 0.91) nor acanthocephalan infection intensity (t = -0.51, *P*-value = 1) had a significant impact on righting response.

Righting response was measured after acclimation (**Figure S4**), and averaged 6.23 seconds ± 10.43 seconds. Genotype significantly impacted TTR (t = 3.65, *P*-value = 0.01), with CC crabs having faster TTR than CT crabs (mean_NH, CC_ = 4.41 seconds ± 6.67 seconds; mean_NH, CT_ = 14.76 seconds ± 14.27 seconds), but the importance of these effects may be impacted by low sample size of CT crabs. No TT crabs were collected from the field for this experiment.

#### Appendix S4: Experiment 2: The role of “heat priming” in shaping thermal tolerance

##### Methods

A total of 24 crabs (carapace width = 47 mm ± 6 mm, weight = 24.18 g ± 8.28 g) were collected from Grays Harbor, WA, USA by Washington Fish and Wildlife and received at Woods Hole Oceanographic Institution on June 22, 2023. Crabs were labelled and sorted by integument color, then held at room temperature (~10-15ºC) for 39 days until genotyping analysis was finished (see *Supergene genotyping*; **Figure S2C**). Crabs were split between four tanks to ensure even distribution of sex, carapace width, integument color, and genotype, and acclimated at 15ºC for one week using a flowing water bath.

##### Oxygen consumption for repeated heat shock experiment

A pilot experiment on the rate of oxygen consumption was additionally used to measure metabolic performance of *C. maenas* to understand how repeated heat shocks influenced physiology. After obtaining righting response measurements on day 4 (after the second heat shock), crabs (n_control_ = 6 and n_treatment_ = 4) weighing ~20g were placed in 90 mm x 50 mm (270 mL) glass respirometry chambers with FireSting oxygen sensor spots (PS-OXSP5). These glass chambers were cleaned with 70% ethanol, covered with dark material to prevent light from entering, and covered with mesh to allow for water exchange. Crabs in the chambers were acclimated for one hour in an aerated tank at the crab’s treatment temperature. Respirometry chambers were refilled with aerated water at the treatment temperature, sealed with parafilm, a glass lid, and a weight to prevent oxygen intrusion, and then placed in a water bath at the treatment temperature. Percent oxygen saturation was measured with a 4-channel FireSting PRO (PyroScience) until readings reached 75% saturation. Respirometry chambers were then flushed and resealed without the crab to measure background respiration for at least 20 minutes. Bare optical fibers for the FireSting devices (PS-SPFIB-BARE) were calibrated using ambient air as the upper limit.

Oxygen consumption rates were compared between control and treatment crabs. To obtain rates for each individual crab, dissolved oxygen (µmol/L) was first calculated from percent air saturation, temperature (ºC), and pressure (mbar) provided by the FireSting and a salinity measurement obtained from each tank with a refractometer. These calculations were performed using the conv_o2 function from the R package respirometry v.2.0.0 (Birk 2023). Linear models were fitted to data when air saturation was between 80-100% saturation. Regression slopes from these models were then obtained for dissolved oxygen (fit from the R package generics v.0.1.3 (Wickham, Kuhn and Vaughan 2022)) and adjusted R-squared values (glance from generics package). Similarly, the last ten minutes of background respiration measurements were used to obtain a “blank” oxygen consumption rate. The slopes from the blank were subtracted from the crab respiration slope to obtain a blank-corrected oxygen consumption slope. A linear model (lm in R) was used to evaluate how treatment impacted oxygen consumption. Genotype effects were not tested due to low sample size.

Oxygen consumption on day 4 was slower for crabs that experienced a prior heat shock (7475.60 nmol/hr ± 4671.50 nmol/hr) than for those that did not (15394.55 nmol/hr ± 8506.37 nmol/hr) (**Figure S6**). However, this trend was not significant (t = -1.68, p = 0.13).

#### Appendix S5: Experiment 3: Changing cold tolerance across populations over time

##### Methods

Crabs (n = 59; carapace width = 47 mm ± 8 mm, weight = 25.26 g ± 15.53 g) from Buzzards Bay, MA, USA were obtained from a bait shop in New Bedford, MA, USA on June 6, 2024. Additional crabs (n = 28; carapace width = 44 mm ± 8 mm, weight = 20.63 g ± 11.63 g) were trapped in Eel Pond, MA, USA between June 7, 2024 and June 15, 2024 under MA Special Permit 174763. Since New Bedford and Eel Pond are both part of Buzzards Bay, MA, USA, crabs from these locations were treated as one MA population (**Figure S2D**). Crabs (n = 98; carapace width = 53 mm ± 4 mm, weight = 32.85 g ± 7.62 g) from Grays Harbor, WA, USA were trapped by Washington Fish and Wildlife on June 26, 2024 (**Figure S2E**). All crabs were sorted by integument color, labelled, and acclimated to ~15ºC in an environmental chamber for two weeks. The MA *C. maenas* were split between six tanks, and the WA crabs were split between eight tanks, with 12-15 crabs per tank. After acclimation, the environmental chamber was set to 0ºC and half of the tanks decreased to a target temperature of 1.5ºC over 12 hours (MA) or 18 hours (WA). The remaining tanks were outfitted with a 300-Watt Deluxe Titanium Heating Tube (Finnex, USA) with an external 1650-Watt temperature controller and ramped down to 5ºC over 10 hours (MA) or 12 hours (WA).

##### Sensitivity test

Sample size decreased over the course of this experiment due to 5-6 crabs per treatment (1-2 crabs per tank) being randomly selected for dissection at each timepoint for future analyses. To account for this, a sensitivity analysis was used to confirm the validity of the modeling results. The same three-step modeling approach used to assess the impact of treatment, time, their interaction, and genotype variables was run using only a subset of the data for which all crabs had data for all timepoints.

Results of the model with all data were compared with results from the subset qualitatively. There were no differences between the most parsimonious models identified by the sensitivity test and the final models for average time-to-right in the MA and WA populations (**Table S8**) and failure-to-right in the MA population. Time did not have a significant impact in the most parsimonious model derived for the sensitivity test for failure-to-right in the WA population. Time was removed from the final model for the full dataset (**Table S8**). Temperature was still significant in the revised model (χ^2^ = 26.57, *P*-value < 0.0001).

#### Supplementary Tables

**Table S1**. PCR primers and amplification methods for SMC gene. A total of 12.5 µL of GoTaq® Master Mix (Promega, USA), 2.5 µL of the forward primer (10 µM), 2.5 µL of the reverse primer (10 µM), and 5.5 µL nuclease-free water was used with 2 µL of DNA for each sample. Additional information on SMC F and R primers can be found in Coyle et al. (2019).

| **Year used** | **Primers** | **Sequence** | **PCR product length (bp)** | **PCR profile** |
| --- | --- | --- | --- | --- |
| 2023  (Experiment 2) | SMC_F  SMC_R | Forward:  5’ - AGCACAGGAAGGCTGTGG - 3’  Reverse:  5’ - ACGAAATCATAAGCCTCTTCACG - 3’ | 126 | 3 min. 94ºC; 17 cycles (30 sec. at 94ºC, 30 sec. at 65ºC [-1ºC per cycle], 1.5 min. at 72ºC). 20 cycles (30 sec. at 94º, 30 sec. at 48ºC 1.5 min. at 72ºC). 10 min. at 72ºC |
| 2024  (Experiments 1 and 3) | SMC_long_F  SMC_long_R | Forward:  5’ - TGATGCTCAGCACAGGAAGG - 3’  Reverse:  5’ - CTTCCATACTTTAACTTCATGAGAACA - 3’ | 690 | 3 min. 95ºC; 35 cycles (30 sec. at 95ºC, 30 sec. at 60ºC, 1 min. at 72ºC). 5 min. at 72ºC |

**Table S2**. Experimental temperatures used for all populations and conditions. Average temperatures are reported for all populations and conditions, except for the second heat shock in experiment 2, where individual tank temperatures are reported due to significant differences in tank temperatures.

| **Experiment** | **Population** | **Condition** | **Temperature (ºC; Mean ± SE)** |
| --- | --- | --- | --- |
| 1 | WA | Heat shock | 30.04 ± 0.30 |
|  | ME | Heat shock | 29.91 ± 0.30 |
| 2 | WA | First Heat Shock | Control: 15.43 ± 0.46  Treatment: 31.55 ± 1.15 |
|  | WA | Second Heat Shock | Control_Tank1_ : 32.14 ± 0.10  Control_Tank 2_ : 32.13 ± 0.07  Treatment_Tank 1_ : 32.09 ± 0.17  Treatment_Tank 2_ : 29.46 ± 0.16 |
| 3 | MA | Cold | 5.39 ± 0.30 |
|  | MA | Colder | 1.66 ± 0.32 |
|  | WA | Cold | 5.20 ± 0.22 |
|  | WA | Colder | 2.14 ± 0.35 |

**Table S3**. Statistical results from testing the importance of demographic variables for average TTR from Experiments 1-3. Likelihood ratio tests were performed with ANOVA to compare the null model with a model testing the influence of a specific variable (ex. integument color). All values are reported after Bonferroni correction.

| **Variable** | **Experiment 1** | | | | **Experiment 2** | | **Experiment 3** | | | |
| --- | --- | --- | --- | --- | --- | --- | --- | --- | --- | --- |
|  | **χ^2^_WA_** | ***P*-value_WA_** | **χ^2^_ME_** | ***P*-value_ME_** | **χ^2^_WA_** | ***P*-value_WA_** | **χ^2^_MA_** | ***P*-value_MA_** | **χ^2^_WA_** | ***P*-value_WA_** |
| Sex | 0.99 | 1 | 0.19 | 1 | 1.35 | 1 | 0.0003 | 1 | 0.27 | 1 |
| Integument Color | 0.68 | 1 | 0.002 | 1 | 2.65 | 1 | 0.17 | 1 | 2.25 | 1 |
| Carapace Width | 0.87 | 1 | 0.35 | 1 | 0.24 | 1 | 0.02 | 1 | 2.05 | 1 |
| Weight | 0.58 | 1 | 0.16 | 1 | 1.05 | 1 | 0.02 | 1 | 0.75 | 1 |
| Missing Fifth Pereopod | 0.14 | 1 | 0.04 | 1 | 5.12 | 0.26 | 0.52 | 1 | 0.43 | 1 |
| Metacercaria Infection Intensity | N/A | N/A | 0.05 | 1 | N/A | N/A | N/A | N/A | N/A | N/A |

**Table S4**. Pairwise statistical test results examining treatment differences at specific timepoints for Experiment 3. All reported statistics are for the 1.5ºC - 5ºC contrast.

| **Hour** | **t_MA, 1.5ºC - 5ºC_** | ***P*-value_MA, 1.5ºC - 5ºC_** | **t_WA, 1.5ºC - 5ºC_** | ***P*-value_WA, 1.5ºC - 5ºC_** |
| --- | --- | --- | --- | --- |
| 0 | 0.27 | 0.79 | 0.49 | 0.62 |
| 4 | 4.44 | < 0.0001 | 5.02 | < 0.0001 |
| 22 | 8.09 | < 0.0001 | 10.15 | < 0.0001 |
| 28 | 9.38 | < 0.0001 | 12.34 | < 0.0001 |
| 46 | 9.50 | < 0.0001 | 12.34 | < 0.0001 |
| 94 | 9.33 | < 0.0001 | 11.85 | < 0.0001 |

**Table S5**. Pairwise statistical test results examining differences in TTR at each timepoint for specific treatments for Experiment 3.

| **Contrast** | **t_MA, 1.5ºC_** | ***P*-value_MA, 1.5ºC_** | **t_MA, 5ºC_** | ***P*-value_MA, 5ºC_** | **t_WA, 1.5ºC_** | ***P*-value_WA, 1.5ºC_** | **t_WA, 5ºC_** | ***P*-value_WA, 5ºC_** |
| --- | --- | --- | --- | --- | --- | --- | --- | --- |
| Hour 0 - Hour 4 | -7.69 | < 0.0001 | -1.80 | 0.47 | -1.22 | 0.83 | -0.85 | 0.96 |
| Hour 0 - Hour 22 | -11.68 | < 0.0001 | -7.49 | < 0.0001 | -16.74 | < 0.0001 | -8.40 | < 0.0001 |
| Hour 0 - Hour 28 | -10.44 | < 0.0001 | -7.34 | < 0.0001 | -17.06 | < 0.0001 | -9.39 | < 0.0001 |
| Hour 0 - Hour 46 | -11.90 | < 0.0001 | -5.49 | < 0.0001 | -18.03 | < 0.0001 | -7.97 | < 0.0001 |
| Hour 0 - Hour 94 | -10.75 | < 0.0001 | -4.13 | 0.0008 | -16.31 | < 0.0001 | -6.16 | < 0.0001 |
| Hour 4 - Hour 22 | -7.49 | < 0.0001 | -5.19 | < 0.0001 | -15.10 | < 0.0001 | -7.38 | < 0.0001 |
| Hour 4 - Hour 28 | -5.59 | < 0.0001 | -5.06 | < 0.0001 | -15.62 | < 0.0001 | -8.443 | < 0.0001 |
| Hour 4 - Hour 46 | -6.96 | < 0.0001 | -3.62 | 0.005 | -16.79 | < 0.0001 | -7.17 | < 0.0001 |
| Hour 4 - Hour 94 | -6.59 | < 0.0001 | -2.62 | 0.10 | -15.22 | < 0.0001 | -5.44 | < 0.0001 |
| Hour 22 - Hour 28 | 2.23 | 0.23 | 0.01 | 1 | -2.05 | 0.32 | -1.56 | 0.63 |
| Hour 22 - Hour 46 | 1.15 | 0.96 | 1.17 | 0.85 | -4.44 | 0.0002 | -1.02 | 0.91 |
| Hour 22 - Hour 94 | 0.71 | 0.98 | 1.61 | 0.59 | -3.90 | 0.002 | 0.39 | 1 |
| Hour 28 - Hour 46 | -1.15 | 0.86 | 1.15 | 0.86 | -2.45 | 0.14 | 0.37 | 1 |
| Hour 28 - Hour 94 | -1.46 | 0.69 | 1.59 | 0.61 | -2.09 | 0.29 | 1.66 | 0.56 |
| Hour 46 - Hour 94 | -0.41 | 1 | 0.55 | 0.99 | 0.19 | 1 | 1.24 | 0.82 |

**Table S6**. Statistical results from the sensitivity analysis testing the importance of variables on average TTR data for the subset of crabs alive throughout Experiment 3. The impact of missing a fifth pereopod was not tested for WA crabs, as no crabs present at the end of the experiment were missing this leg. Only full genotype was tested due to TT crabs not being present at the end of the MA experiment and CC crabs not being present at the end of the WA experiment. Likelihood ratio tests were performed with ANOVA to compare the null model with a model testing the influence of a specific variable (ex. integument color). All values are reported after Bonferroni correction.

| **Variable** | **χ^2^_MA_** | ***P*-value_MA_** | **χ^2^_WA_** | ***P*-value_WA_** |
| --- | --- | --- | --- | --- |
| Sex | 0.21 | 1 | 0.60 | 1 |
| Integument Color | 0.34 | 1 | 0.27 | 1 |
| Carapace Width | 0.01 | 1 | 0.18 | 1 |
| Weight | 0.004 | 1 | 1.66 | 1 |
| Missing Fifth Pereopod | 0.75 | 1 | N/A | N/A |
| Temperature | 48.98 | < 0.0001 | 77.30 | < 0.0001 |
| Time | 62.64 | < 0.0001 | 146.38 | < 0.0001 |
| Temperature:Time | 26.35 | < 0.0001 | 34.94 | < 0.0001 |
| Genotype | 0.30 | 1 | 0.09 | 1 |

**Table S7**. Statistical results from testing the importance of demographic variables for failure to right in Experiment 3. Likelihood ratio tests were performed with ANOVA to compare the null model with a model testing the influence of a specific variable (ex. integument color). All values are reported after Bonferroni correction.

| **Variable** | **χ^2^_MA_** | ***P*-value_MA_** | **χ^2^_WA_** | ***P*-value_WA_** |
| --- | --- | --- | --- | --- |
| Sex | 0.41 | 1 | 0.54 | 1 |
| Integument Color | 0.19 | 1 | 0.01 | 1 |
| Carapace Width | 0.58 | 1 | 2.50 | 1 |
| Weight | 0.68 | 1 | 1.60 | 1 |
| Missing Fifth Pereopod | 0.03 | 1 | 0.15 | 1 |

**Table S8**. Statistical results from the sensitivity analysis testing the importance of variables on failure to right data for the subset of crabs alive throughout Experiment 3. The impact of missing a fifth pereopod was not tested for WA crabs, as no crabs present at the end of the experiment were missing this leg. Only full genotype was tested due to TT crabs not being present at the end of the MA experiment and CC crabs not being present at the end of the WA experiment. Likelihood ratio tests were performed with ANOVA to compare the null model with a model testing the influence of a specific variable (ex. integument color). All values are reported after Bonferroni correction.

| **Variable** | **χ^2^_MA_** | ***P*-value_MA_** | **χ^2^_WA_** | ***P*-value_WA_** |
| --- | --- | --- | --- | --- |
| Sex | 0.0002 | 1 | 3.05 | 0.65 |
| Integument Color | 1.35 | 1 | 0.01 | 1 |
| Carapace Width | 0.02 | 1 | 0.23 | 1 |
| Weight | 0.15 | 1 | 1.17 | 1 |
| Missing Fifth Pereopod | 1.00 | 1 | N/A | N/A |
| Temperature | 24.33 | < 0.0001 | 17.11 | 0.0002 |
| Time | 20.81 | 0.0003 | 7.43 | 0.19 |
| Temperature:Time | 0 | 1 | 1.76 x 10^-7^ | 1 |
| Genotype | 3.72 | 0.48 | 2.25 | 1 |

#### Supplementary Figure Captions

**Figure S1**. Example gel showing genotyping with AluI restriction digest. **1**) Undigested PCR product (690 bp). A faint band is also present between 300-400 bp that is likely a result of non-target amplification. Presence of this non-diagnostic band can vary across CC, CT, and TT samples. **2**) CC band (389 bp). **3**) CT bands (477 bp and 389 bp). **4**) TT band (477 bp). Primer-dimer bands for all samples and the non-diagnostic 88 bp bands for CC and CT genotypes are present in all digested samples. Shown with a 100 bp ladder for reference.

**Figure S2**. Pie charts for supergene genotypes in **A**) WA and **B**) ME populations used in experiment 1, **C**) WA population used in experiment 2, and **D**) MA and **E**) WA populations used in experiment 3. Percentages for each genotype are presented.

**Figure S3**. Experimental temperature conditions for **A**) WA (purple) and **B**) ME (green) experiments; HOBO loggers were added to the tanks after the heat ramp had started in **A**).

**Figure S4**. Righting response in NH *C. maenas* by genotype. CT crabs had significantly slower righting response at room temperature than CC crabs (t = 10.84, P-value = 8.96 x 10^-3^).

**Figure S5**. **A**) Water temperatures for Experiment 2. Vertical lines indicate when TTR was measured (day 0, day 1, or day 4). Dashed grey horizontal lines indicate temperature conditions for the control crabs averaged across two replicate tanks, while dashed red horizontal lines indicate temperature conditions for treatment crabs averaged across two replicate tanks. Standard error during the second heat stress is larger for the treatment crabs due to slight differences in temperature conditions between replicate tanks. **B**) Tank-specific temperatures during the second heat stress. One treatment tank had a significantly lower temperature than the other three tanks. In all panels, red indicates treatment tanks and grey indicates control tanks.

**Figure S6**. Oxygen consumption (nmol/hr) after the day 4 heat shock. Crabs that did not experience a prior heat shock are in grey, while those that did are in red. Different shapes indicate crab genotype. There was no significant impact of prior heat shock exposure on oxygen consumption.

**Figure S7**. Temperature conditions for **A**) MA and **B**) WA *C. maenas* used in experiment 3. In both panels, average temperatures experienced by crabs across replicate tanks in cold (light blue; MA = 5.39ºC ± 0.30ºC; WA = 5.20ºC ± 0.22ºC) and colder (dark blue; MA = 1.66ºC ± 0.32ºC; WA = 2.14ºC ± 0.35ºC)) treatments are indicated by the solid lines. Ribbons depict standard errors. TTR sampling points are indicated by vertical lines. Experiments were conducted in 2024.

**Figure S8**. Righting responses for crabs that were alive throughout the duration of Experiment 3. Binary righting response for **A**) MA and **B**) WA crabs kept at 1.5ºC. Average time-to-right for **C**) MA and **D**) WA kept at 5ºC. Hour 0 is not shown as it is before the cold exposures began.
